# Single Nucleotide Mapping of the Locally Accessible Trait Space in Yeast Reveals Pareto Fronts that Constrain Initial Adaptation

**DOI:** 10.1101/593947

**Authors:** Yuping Li, Dmitri A. Petrov, Gavin Sherlock

## Abstract

Tradeoffs constrain the improvement of performance of multiple traits simultaneously. Such tradeoffs define Pareto fronts, which represent a set of optimal individuals that cannot be improved in any one trait without reducing performance in another. Surprisingly, experimental evolution often yields genotypes with improved performance in all measured traits, perhaps indicating an absence of tradeoffs at least in the short-term. Here we densely sample adaptive mutations in *S. cerevisiae* to ask whether first-step adaptive mutations result in tradeoffs during the growth cycle. We isolated thousands of adaptive clones evolved under carefully chosen conditions and quantified their performances in each part of the growth cycle. We too find that some first-step adaptive mutations can improve all traits to a modest extent. However, our dense sampling allowed us to identify tradeoffs and establish the existence of Pareto fronts between fermentation and respiration, and between respiration and stationary phases. Moreover, we establish that no single mutation in the ancestral genome can circumvent the detected tradeoffs. Finally, we sequenced hundreds of these adaptive clones, revealing novel targets of adaptation and defining the genetic basis of the identified tradeoffs.

## Introduction

That gain must ultimately be associated with some cost is a fundamental premise in fields spanning economics, engineering, and biology. Biology in particular has a rich tradition of both alluding to and attempting to define tradeoffs: here tradeoffs imply that a part of trait space is not accessible by evolution, such that, within a defined period of time, a lineage cannot evolve improved performance of two or more traits simultaneously above some threshold. Such evolutionary tradeoffs have been suggested by various biological phenomena - for instance, organisms with high fecundity tend to have a short lifespan ^1–3^ and organisms with large eggs tend to lay fewer of them ^4,5^.

Despite the plethora of such examples of negative correlations between specific traits, such correlations alone are insufficient to demonstrate the existence of tradeoffs. Indeed, many alternative explanations exist. For instance, consider an environment in which only one trait is under selection while a second is not. Over evolutionary time, performance in the first trait is likely to increase while performance of the second is likely to decrease due to the accumulation of damaging mutations in the absence of purifying selection^6,7^. At the same time, a reciprocal relationship may be observed in an alternative environment if the second trait is subject to selection and the first one is not. This will lead to a negative correlation between performances of the two traits. However, it is entirely possible that mutations that improve both traits do exist, but they are not particularly common and not particularly advantageous in either of the environments. Additional explanations, such as sexual selection driving some traits to seemingly suboptimal states^8^, or current selective pressures not corresponding to the way natural selection acted in the past might also lead to negative correlations among traits in the absence of tradeoffs. In short, negative correlation in performance between two traits is expected in the presence of tradeoffs but in and of itself is not sufficiently strong evidence for the existence of tradeoffs.

Consider an organism with two traits under selection (Fig. 1a): its trait-fitness space is two-dimensional, with each axis representing performance for one of the traits. If a tradeoff exists between the two traits, for every biologically possible value of trait 1, the best value for trait 2 performance will be constrained by trait 1, generating a Pareto optimality front (or Pareto front)^9^. Such a Pareto front not only represents the set of optimal trait combinations, but also separates the “accessible” from the “inaccessible” trait space. For individuals on the Pareto front (green dots in Fig. 1a), the existence of tradeoffs can be demonstrated straightforwardly: increasing the performance for one trait will inevitably decrease performance for another. By contrast, individuals behind the Pareto front (the black dot in Fig. 1a) are able to improve performance in both traits simultaneously. It is generally assumed that organisms should be located on or near a “long-term” Pareto front as they are products of very long term evolution^1,2,5,9–12^. Surprisingly, results from experimental evolution often demonstrate the improvement of multiple traits simultaneously, suggesting that at least for the conditions and traits tested, the ancestor does not lie on a Pareto front^13–19^. However, it is important to appreciate that it is possible for an individual to be on a higher dimensional Pareto front, defined by multiple traits, but when measuring only a subset of the traits, the organism will appear to be behind the front (Fig. 1b). In this case, improvement in performance in the subset of traits must come at the cost of performance in the additional, unmeasured, traits that contribute to the higher dimensional front.

**Figure 1:**
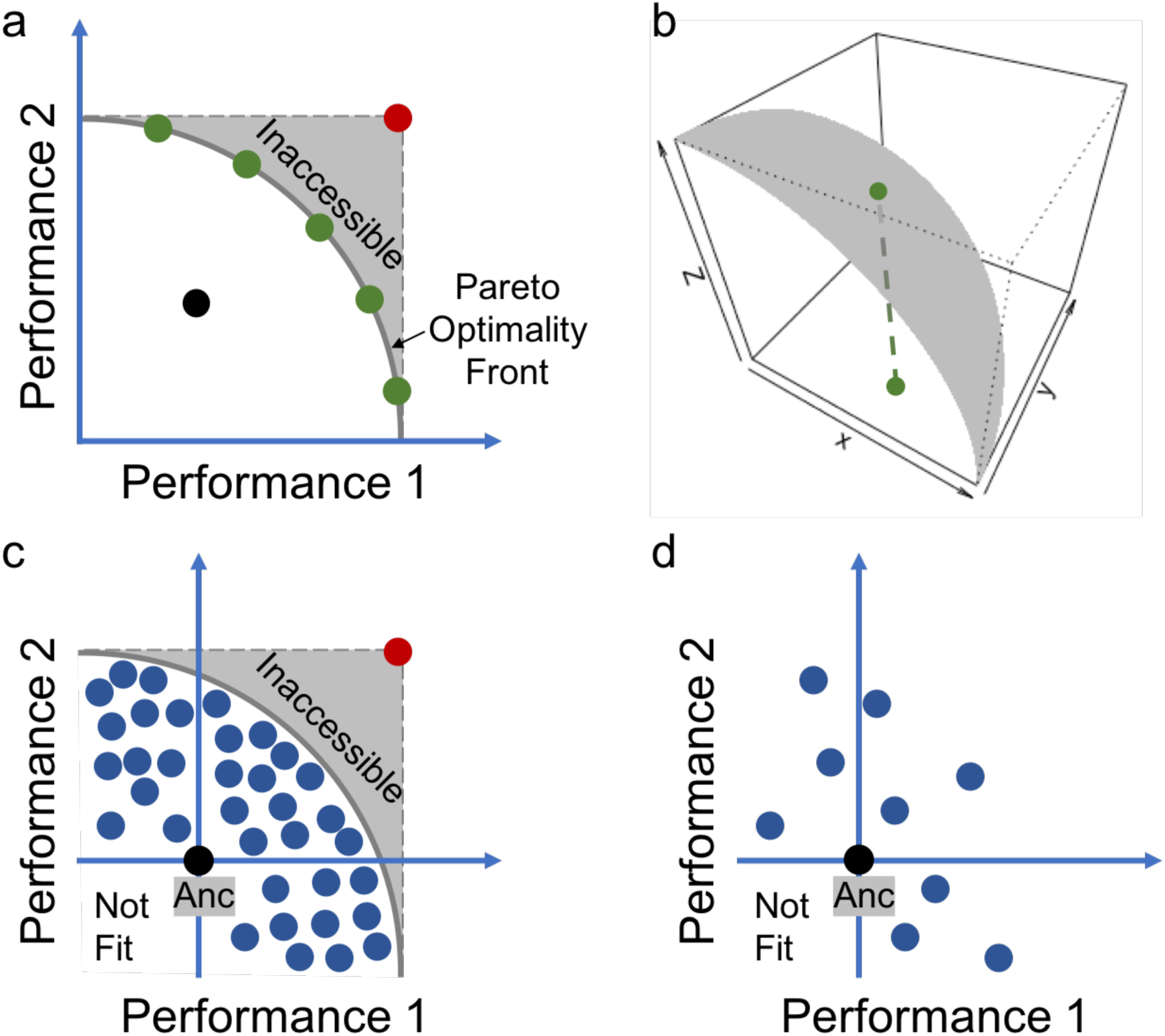
Evolutionary constraints in trait-performance space. **a,** The Pareto optimality front separates the evolutionary accessible (white space) from the inaccessible space (shaded space). The red dot represents mutants that maximize both traits simultaneously. When organisms are on the Pareto optimality front (green dots), increasing the performance for one trait decreases the performance for the other. By contrast, when organisms are behind the Pareto front (black dot), organisms can improve the performance of both traits until the front is reached. **b,** An organism on a three-dimensional Pareto surface (green dot) appears to be sub-optimal when it is projected onto a two-dimensional space. **c-d,** When the ancestor (Anc) is behind the Pareto front, many individuals occupying different parts of the trait space (**c**) are required to characterize the Pareto front. By contrast, too few individuals (**d**) are insufficient to delineate the front.

The Pareto front is typically thought of as being defined by physical, structural, or physiological constraints. However, the Pareto front may also be defined by genetic constraints, such that the space above the front might be locally inaccessible in the short-term due to the rarity of specific genetic changes required to reach that region of trait space. For example, if the “inaccessible” part of trait space requires the system to move through a fitness valley the system might remain at the ‘Pareto front’ at least in the short term. The transition into the locally inaccessible part of the space would then be seen as a true evolutionary innovation that shifts the Pareto front to a new location. The Pareto front is thus defined both by the timescale of evolution and the physiological or structural relationships among the traits.

To explore whether even the first step of adaptation can reveal evolutionary constraints in the form of Pareto fronts, one needs to sample a large number of adaptive mutants selected for multiple traits under a range of conditions and then precisely measure their performance along each trait axis (Fig. 1c,d). Pareto fronts, if present, can then be inferred by an absence of mutants able to maximize both traits simultaneously (the large red dot in Fig. 1a,c). If the first step mutations can reach the short-term Pareto optimality front and if the density of sampling is such that any adaptive single-step mutant that would land beyond the defined front would have been detected with high likelihood, then a short-term Pareto front will have been demonstrated.

Here we set out to investigate the existence of Pareto fronts among multiple traits, by evolving barcoded yeast populations under a number of carefully chosen conditions, selecting for improved performance in different phases of the yeast growth cycle, including fermentation, respiration, and stationary phases. We isolated ∼500 independent adaptive clones most of which carry a single beneficial mutation. We found that a number of adaptive clones improved all three measured performances to a modest extent without apparent tradeoffs, indicating that the ancestor cannot be located on a Pareto front for the measured traits. However, no adaptive clones were able to maximize performance in some pairs of traits. We were able to delineate apparent short-term Pareto fronts between fermentation and respiration as well as between respiration and stationary phases, but not between fermentation and stationary phase performances. Importantly, due to a large number of sampled and tested clones we could assert that no single point mutation in the yeast genome can improve the performance substantially beyond either of the two defined Pareto fronts. Finally, by sequencing hundreds of adaptive clones, we identified the genetic basis underlying the identified tradeoffs and revealed novel targets of adaptation.

## Results

### Experimental System and Isolation of Evolved Clones

When yeast cells grow in conditions with a fermentable carbon source, such as glucose used in this study, they go through a sequence of growth phases: (i) lag phase, where cells acclimate to the medium, with no cell division; (ii) fermentation, where cells divide exponentially by converting glucose into ethanol; (iii) respiration, where glucose is exhausted and cells divide slowly by consuming the ethanol produced during fermentation; and (iv) stationary/starvation phase, where cells cease growth because readily-available carbon has been depleted from the medium (Fig. 2a).

**Figure 2:**
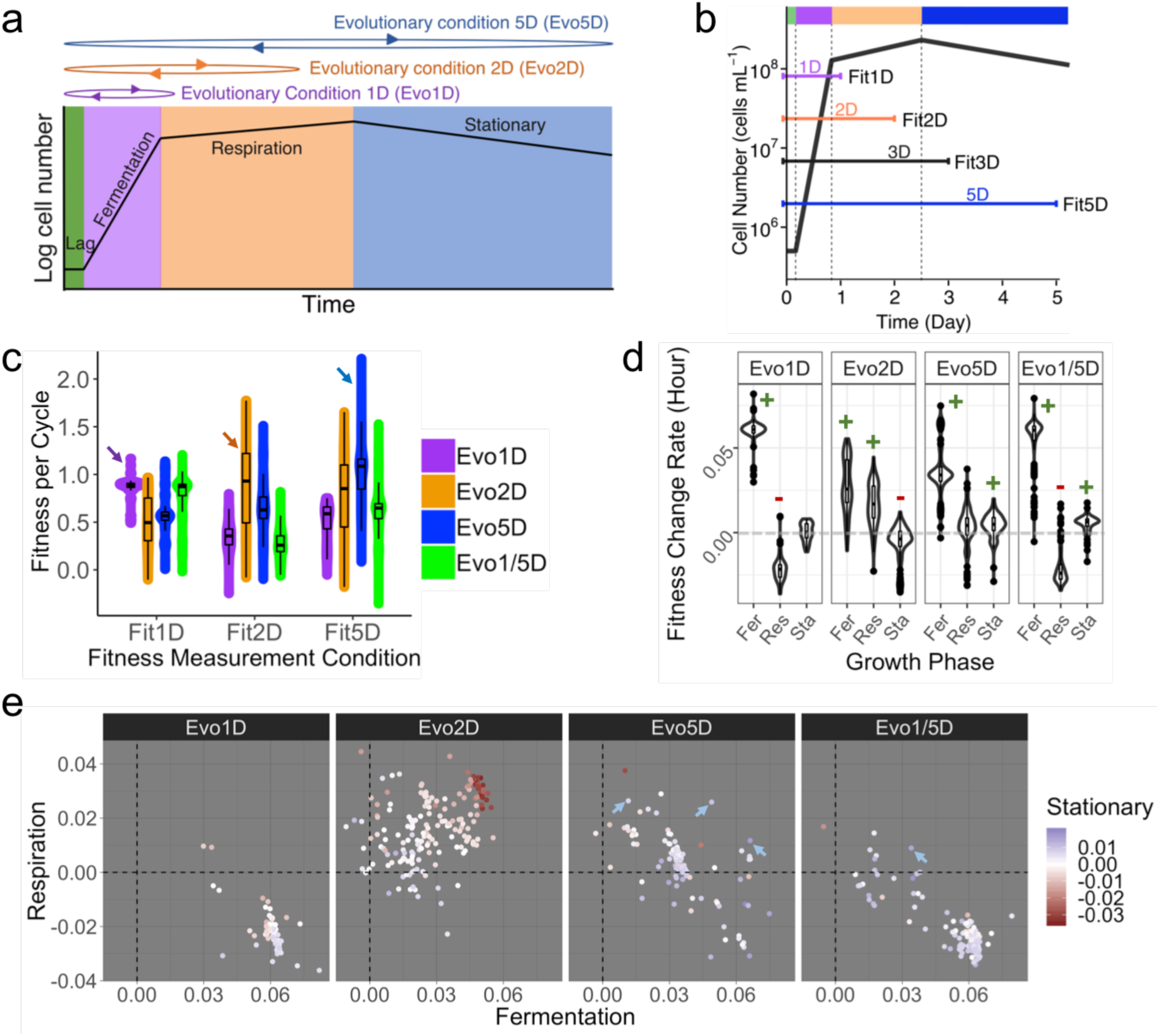
Experimental design and the observation of local adaptation and tradeoffs. **a,** Three chosen evolutionary conditions span different phases of the yeast growth cycle. Clones were also evolved in a 1-day/5-day alternating condition (Evo1/5D). **b,** Fitness measurement conditions designed to quantify fermentation, respiration and stationary performances (fitness change per hour) of each clone. Dashed vertical lines separate different growth phases, colored as (**a**). **c,** Fitness measurements of adaptive clones, grouped by their “home” evolutionary condition, in “home” and “away” conditions. Arrows point to adaptive clones measured in their “home” condition. **d,** Adaptive clones’ fermentation, respiration and stationary performances grouped by their evolutionary condition. +/-indicates increased/decreased performance compared to the ancestor. **e,** Clones are separated by their evolutionary condition and colored by their stationary phase performance. Each dot represents a clone. Note that some blue colored clones from Evo5D and Evo1/5D (pointed by arrows) improve performances in all three growth phases.

To isolate adaptive clones with improved performances in fermentation, respiration, and/or stationary phase (or combinations thereof) we propagated barcoded haploid yeast populations under four serial transfer conditions having differing cycle lengths: 1) 1-day (referred to as Evo1D below) including 4h lag, 16h fermentation, and 4h respiration; 2) 2-day (Evo2D, conducted in Levy, Blundell *et al*^20^) including additional 24h respiration; 3) 5-day (Evo5D) including a further 12h respiration and 60h stationary phase; and 4) alternating 1-day and 5-day transfer (Evo1/5D) (Fig. 2a). We used barcode trajectories to determine that cell cultures in cycle 11 contained a high proportion of diverse adaptive clones. Furthermore, our previous analysis indicated that at this time point most adaptive clones would contain only a single adaptive mutation^20^. Subsequent sequencing of individual clones (Venkataram, Dunn *et al*^21^ and see below) confirmed this supposition.

We isolated clones from cycle 11 for subsequent analysis. Specifically, from Evo1D, Evo2D, Evo5D, and Evo1/5D we isolated respectively 120, 3048 (isolated in Venkataram, Dunn *et al*^21^), 157, and 384 distinct evolved clones carrying unique barcodes. We previously found that ∼50% of clones isolated from Evo2D had self-diploidized during the course of evolution^21^ and were beneficial across all fitness measurement conditions^22^. We therefore assayed the ploidy of newly isolated clones, and observed 43%, 45%, and 14% diploids among clones isolated from Evo1D, Evo5D, and Evo1/5D, respectively.

We measured the fitness of all isolated clones in 1-day (Fit1D), 2-day (Fit2D), 3-day (Fit3D) and 5-day (Fit5D) serial transfer conditions (Fig. 2b, clones from Evo2D were measured in Li, Venkataram *et al*^22^) using the method developed in Venkataram, Dunn *et al*^21^. For each clone, we therefore have its fitness in the “home” condition (except for Evo1/5D clones), as well as the “away” conditions. Note that one condition (Fit3D) was not used as an evolutionary condition but instead was important for evaluating stationary phase performance. Below we use these values to investigate patterns of local adaptation and to estimate performance of each clone in fermentation, respiration, and stationary phases. Using the fitness and ploidy measurements, we identified 66, 144, 58, and 132 adaptive haploids and 4, 40, 57, and 6 high-fitness diploids (assumed to have additional beneficial mutations besides diploidy) from Evo1D, Evo2D, Evo5D, and Evo1/5D, respectively. We refer to these adaptive haploids and high-fitness diploids collectively as adaptive clones.

### Local Adaptation Results from Performance Differences in Different Growth Phases

We observed a large range of fitness both in the “home” and “away” environments (Fig. 2c). For example, the fitness of all adaptive clones varied from −0.35 to +2.2 per growth cycle in Fit5D, suggesting multiple adaptive strategies and targets of adaptation among these clones. While only 4.5% of the adaptive clones are maladaptive in any away condition, we do find that in general, adaptive clones exhibit evidence of local adaptation. Specifically, for each fitness remeasurement condition, both the average and the highest fitness of clones evolved in the home condition (indicated by arrows) are greater than those of clones evolved in the away conditions. Nonetheless, under a given fitness measurement condition, not all “home” clones are more fit than all “away” clones.

We further used our combined fitness data to determine the *performance* of individual clones in three of the phases in the growth cycle: fermentation, respiration, and stationary phase (Fig. 2d). Here, we define performance as the increase in fitness, *per hour*, for a given growth phase; our previous study demonstrated that the overall fitness scales linearly with the amount of time spent in each of the growth phases^22^. The slope of the relationship between the relative fitness of a clone and the length of a particular growth phase (measured as fitness change per hour) can thus be used as a measure of clone performance in that phase. For instance, as the clones spend 24 extra hours in respiration during every cycle when growing under Fit2D compared to Fit1D we can calculate respiration performance by subtracting relative fitness of each clone in Fit1D from that in Fit2D and then dividing by 24 hours. Similarly, we calculated the fermentation and stationary performances (Supplementary Information section 6).

We compared these three performances for clones evolved in all four conditions. Overall, while clones from each condition often revealed specific and consistent patterns of apparent tradeoffs, the tradeoffs observed were not necessarily shared across all conditions (Fig. 2d,e). For example, we previously found that most adaptive clones from Evo2D have improved performance in both fermentation and respiration, but decreased performance in stationary phase^22^. By contrast, adaptive clones from Evo1D have improved performance in fermentation, yet decreased performance in respiration and nearly unchanged performance in stationary phase. Most adaptive clones from Evo5D exhibit yet a different pattern -- improved performance in both fermentation and stationary phases but their performance in respiration on average is largely unchanged. Finally, adaptive clones from Evo1/5D have improved fermentation and stationary phase performance and generally decreased respiration performance. Overall, we found adaptive clones that improved every pair of fermentation, respiration, and stationary phase performances, as well as some that showed improved performance across all three (indicated by arrows in Fig. 2e), suggesting that the ancestor is behind any potential Pareto front for these three performances.

### The Genetic Basis of Adaptation and Tradeoffs

We determined the genetic basis of adaptation by genome-wide sequencing of 47, 67, and 85 adaptive clones from Evo1D, Evo5D, and Evo1/5D respectively. Putative adaptive mutations were successfully identified in 35 (74%), 66 (98%), and 81 (95%) of these clones. The identity of 125 adaptive mutants from Evo2D was determined previously^21,22^. Many genes or pathways were recurrently mutated in our adaptive clones – in such cases we can be confident that these mutations are indeed adaptive. Specifically, out of the 182 adaptive clones in which we identified putative adaptive mutations, 118 (∼65%) harbor mutations in genes/pathways hit in multiple clones (Table S3). Furthermore, 79 of them harbor mutations in genes/pathways independently hit five or more times (Table 1).

**Table 1:**
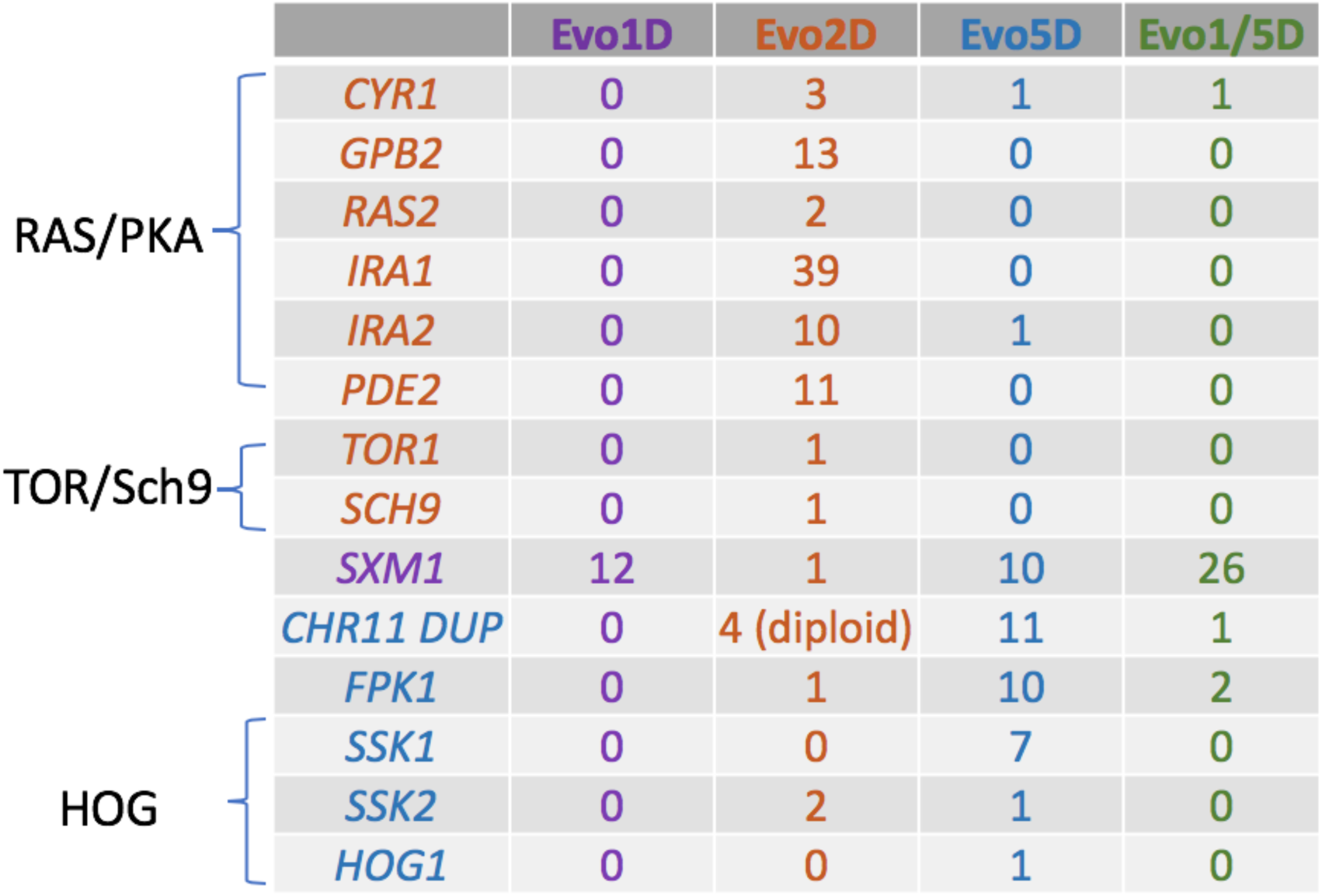
Genetic basis of adaptation and tradeoffs. The number of clones carrying recurrent mutations within genes or pathways. These genes/pathways were independently mutated more than four times. Genes in the same pathway are grouped by the large parenthesis on the left.

In general, within each evolutionary condition, beneficial mutations were limited to a small number of genes that serve similar biological functions. At the same time, across evolutionary conditions, beneficial mutations tend to differ in their genetic bases (Table 1). For instance, we previously reported that the majority of adaptive mutants for Evo2D upregulated the *RAS/PKA* and *TOR/Sch9* nutrient sensing pathways^21^, but we rarely recovered adaptive mutations in these pathways from the other evolutionary conditions. By contrast, loss of function in *SXM1* (a nuclear transport factor interacting with the nuclear pore complex^23^) was the prevalent cause of adaptation in Evo1D. While *SXM1* mutants were also observed in Evo5D, they were not the predominant mutant class. Instead, a wide variety of mutations were observed among Evo5D adaptive clones, including (i) 11 duplications of chromosome 11 (*Chr11Dup*), (ii) 10 independent loss of function mutations in *FPK1*, and (iii) 9 mutations in three components of the high-osmolarity glycerol (*HOG*) response pathway: *SSK1, SSK2,* and *HOG1*. Given that Evo5D contains a long period of starvation, observation of *Chr11* aneuploidy is consistent with previous findings that aneuploidies can improve survival under extremely stressful conditions^124–26^, although the underlying mechanism is unknown. *FPK1* (a flippase activator) has been previously shown to increase viability in stationary phase^27^, which we experimentally confirmed (Table S4). The genetic bases of adaptation among Evo1/5D clones were similar to those for Evo5D clones, with mutations in *SXM1* and *FPK1* as well as duplication of *Chr11*.

Next, we examined the relationship between the identified genetic basis of adaptation and the resulting increases/decreases in performance (Fig. 3a-c). As stated above, in this study “performance” represents fitness change per hour in a particular growth phase rather than measurements of physiological traits (e.g. growth rate) as it is commonly used. The *SXM1* mutants, predominant in Evo1D, have among the highest observed fermentation performances, at >6% per hour (giving >96% fitness advantage over the ancestor over the full 16-hour period of fermentation in our conditions). This likely explains why nutrient-sensing pathway mutants, which have lower fermentation performance, and are common in Evo2D, were not observed in Evo1D. However, the high fermentation performances of *SXM1* mutants come at a cost of reduced respiration performance (negative 2-3% per hour). This likely explains their near absence in Evo2D given that the Evo2D condition contains a long period of respiration. Similarly, the most prevalent Evo2D *RAS/PKA* nutrient-sensing pathway mutants with the highest respiration performance tradeoff strongly in stationary phase^22^, explaining why they were not observed in Evo5D. Finally, clones that are common in Evo5D, which contains all phases of the growth cycle, are the least likely to show decreased performance in any phases of the growth cycle. Indeed, Evo5D specific mutations, such as *Chr11* duplication and *SSK1* mutation, show no obvious tradeoffs, but rather modest improvements in one or more performances (Fig. S1). Interestingly, Evo5D clones also include *SXM1* mutants that show increased performance only in fermentation with decreased performance in respiration and little change in stationary phase. In this case, their strong improvement in fermentation and lack of tradeoff in stationary phase appears to compensate enough for their reduced fitness in respiration.

**Figure 3:**
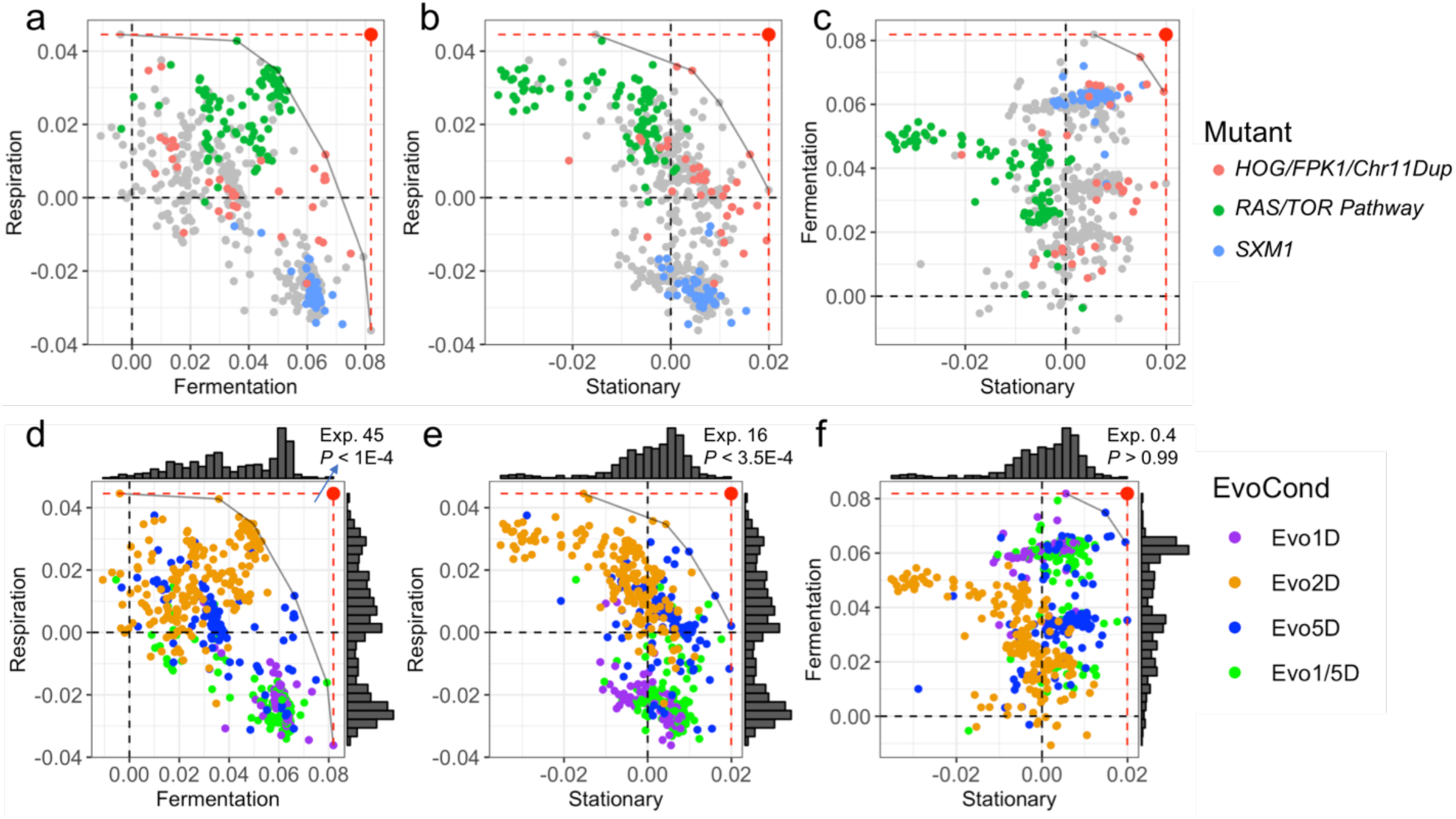
Mapping of the evolutionarily accessible trait space. For each pair of performances (fitness change per hour in each growth phase), adaptive clones are plotted and colored by either their molecular basis (**a-c**), or their evolutionary conditions (**d-f**). Each dot represents a clone. The large red dots represent the optimum phenotypes, achieving the upper limits (dashed lines) of each pair of performances. The grey curves, defined by the convex hull algorithm, represent putative Pareto optimality fronts. **d-f,** Histograms on the side represent the density distribution of each trait’s performance. Based on the null distribution, the number of clones expected to be observed (Exp.) in the empty space between the putative front and the optimal type (the large red dot) is reported, along with the p-value of not observing any clone in this empty space.

In summary, adaptation under these conditions is idiosyncratic yet predictable: the genetic basis of adaptation under a particular evolutionary condition tends to target a narrow, recurrent and thus *a posteriori* predictable set of genes. However, these gene targets are not shared across all environments, meaning that adaptation across conditions often relies on entirely different genetic pathways. This idiosyncratic nature explains the specific patterns of performances across conditions (Fig. 2d,e). While we do detect clones that increase all performances, clones that perform best in any one growth phase tend to tradeoff in performance in some other growth phase(s). This hints at the existence of evolutionary constraints, preventing the emergence of adaptive clones that simultaneously *maximize* performance in all growth phases.

### Identification of Evolutionary Constraints and Delineation of Pareto Fronts

We observed an absence of clones near the upper limits of either both fermentation and respiration performances, or both respiration and stationary performances (the large red dot in Fig. 3a,d and 3b,e). Thus, there is at least the appearance of an empty space in the upper right corner, where these pairs of performances would be maximized. We used the convex hull algorithm to delineate potential Pareto fronts that separate the short-term evolutionarily-accessible space from the empty, putatively short-term inaccessible space above the front (grey curves in Fig. 3).

We first tested whether, given the marginal distributions of trait performances, the absence of clones at the top right of those plots is statistically unexpected (Supplementary Information section 10). Under a null hypothesis of independence of performances, the observation of no clones beyond these putative fronts is indeed unexpected (*P* < 1E-4 for fermentation and respiration phases and 3.5E-4 for respiration and stationary phases, respectively; Fig. 3d,e and S2a,b). By contrast, there is no unexpected lack of clones close to the upper limits of both fermentation and stationary performances (Fig. 3f and S2c; *P* > 0.99).

### The Mutational Target Size of the Optimal Types is Smaller Than A Single Nucleotide

To further explore the absence of clones beyond the putative Pareto fronts, we determined the target size for possible single-step mutations that would give rise to the maximum performances for fermentation and respiration, or respiration and stationary phase (marked by the large red dot in Fig. 3a,d and 3b,e). Mutants that could maximize two traits simultaneously would be more fit than the observed mutants at least in some evolutionary conditions; thus, based on this increased fitness, such mutants, should they arise at a similar rate as the observed mutants, should be sampled frequently in those conditions. For example, mutants that improve fermentation and respiration simultaneously beyond the putative front should have a higher fitness than most of sampled clones in Evo2D (Fig. S3a), as clones in this condition experience only fermentation and respiration. Likewise, clones that improve respiration and stationary phase beyond the putative front should have a high fitness in Evo5D (Fig. S3b), given that the majority of clones with high respiration or stationary performance have a positive fermentation performance as well. The fact that we didn’t observe any clones beyond the putative fronts suggests that the genomic mutational target size towards such extremely fit mutants located beyond the putative Pareto fronts must be smaller than that for the observed mutants.

Next, we used a mathematical model to quantitatively assess the probability of sampling a single-step mutation with a given selection coefficient *s* (Supplementary Information section 11). Several factors determine the probability of sampling such a single-step mutation: the rate at which a mutation occurs, the probability of such a mutation surviving random drift and establishing in the population (∼ proportional to *s*), and the exponential division rate after the mutation establishes (its cell number roughly reaches *e*^*(s*t)*, with *t* generations between establishment and sampling). With mutations entering the population at a fixed rate, the more fit a mutant is (the larger *s* is), the more likely the mutant establishes in the population, the faster the mutant divides and eventually the higher frequency the mutant reaches by the sampling time.

First, consider a gene with the same target size for adaptive mutations as *IRA1* (which were observed 39 times after sampling at cycle 11 of Evo2D ^21,22^), but whose mutation results in a fitness benefit at the hypothetical optimal type, with maximal fermentation and respiration (the red dot in Fig. 3a,d). Such a hypothetical mutant would have a fitness of ∼2.56 per cycle in Evo2D, compared to ∼1.64 per cycle for *IRA1*-nonsense mutations. If such a hypothetical gene exists, we would expect to observe mutations in this gene ∼25,000 times more frequently than we observed mutations in *IRA1* in Evo2D. Thus, it is exceptionally unlikely that such a gene with a similar target size to *IRA1* does exist. Furthermore, if the target size for such a gene is just a single base pair, our mathematical model suggests that we would expect to see such a mutation 84 to 99 percent of the time in our evolution experiments (Supplementary Information section 11). Thus, we believe it is unlikely that there is even a single site in the genome of the ancestral strain that can be mutated to provide such a high fitness.

Similarly, the hypothetical optimal type which maximizes the respiration and stationary phase performances would have a fitness benefit ∼2.98 per cycle in Evo5D (represented by the red dot in Fig. 3b,e) (assuming a fermentation performance of zero). If a single site (1bp) can be mutated to this hypothetical optimal type, we would expect to sample such a mutant 88 to 98 percent of the time in Evo5D experiments. Thus, there is likely no single-step mutation in the ancestral yeast genome that can simultaneously maximize either both fermentation and respiration, or both respiration and stationary performances to their highest observed levels.

## Discussion

### A Large Number of Diversely Selected Adaptive Clones Is Needed to Delineate Pareto Fronts

Despite the fact that tradeoffs have been widely assumed in studies of evolution, it is extremely challenging to formally establish the existence of tradeoffs. Here, by sampling a large number of adaptive clones from a range of evolutionary conditions, and measuring their performance in three different traits, we were able to demonstrate the existence of Pareto fronts between fermentation and respiration, and between respiration and stationary phase performances. Furthermore, we were able to show that the ancestor must be behind these fronts, because for both pairs of traits there were clones that were able to improve performance in both traits simultaneously; indeed, some clones were able to improve performance in all three traits.

If the ancestor was on a front delineated by two traits, characterization of the front using experimental evolution would be straightforward, because no adaptive clones could improve both traits simultaneously – indeed, by definition, improvement of performance in one trait would lead to a loss of performance in the other. However, because the ancestor lies behind the fronts we identified, only by mapping a very large number of adaptive clones whose performances span the trait space could we map the Pareto fronts. By randomly subsampling our data, we estimated that ∼100-200 independent adaptive mutants are required to detect the Pareto fronts in our experiment (Supplementary Information section 10). Furthermore, given that clones isolated from a particular evolutionary condition, e.g. Evo1D, tend to occupy a specific part of the trait space, clones from Evo1D, Evo2D, and Evo5D together were required to detect the Pareto fronts.

Finally, having such a large number of adaptive clones enabled us to show that for both of the identified Pareto fronts there is no single mutation that can occur in the genome of the ancestral strain that would enable the strain to maximize performance in both traits. These fronts therefore constrain the evolutionarily accessible space over short timescales.

### No Observed Pareto Front between Fermentation and Stationary Phase

We were unable to identify a Pareto front between fermentation and stationary phase performances, suggesting either an absence of tradeoffs between these two traits or that single-step mutations provide insufficient performance improvement to reach a hypothetical Pareto front between these two traits. However, this may also be due to experimental limitations: specifically, clones selected under Evo5D experienced both fermentation and respiration prior to stationary phase. Thus, it is entirely possible that the maximum stationary phase performance is larger than we observed, if clones with such a large stationary phase performance tradeoff strongly in fermentation or respiration. A longer stationary phase, e.g. a 10-day serial transfer, may help select for such mutants and define a Pareto front between fermentation and stationary phase performances should one exist. Additionally, evolution in a non-fermentable carbon source followed by a long stationary phase may also enable selection of clones with high stationary phase performance that tradeoff strongly in fermentation.

### The Shape of Pareto Fronts and Nature of Tradeoffs

Levin (1962)^28^ suggested that the geometry of Pareto fronts will affect an organism’s evolvability, and whether generalists or specialists will tend to evolve. For instance, a convex-shaped front allows for better evolvability and produces different optimal types based on the particular evolutionary condition, allowing for local adaptation (Fig. 4a). By contrast, a concave-shaped front leads to less evolvability, because regardless of the importance of performance in each trait, one of the two most specialized types will always be the most fit (Fig. 4b).

**Figure 4:**
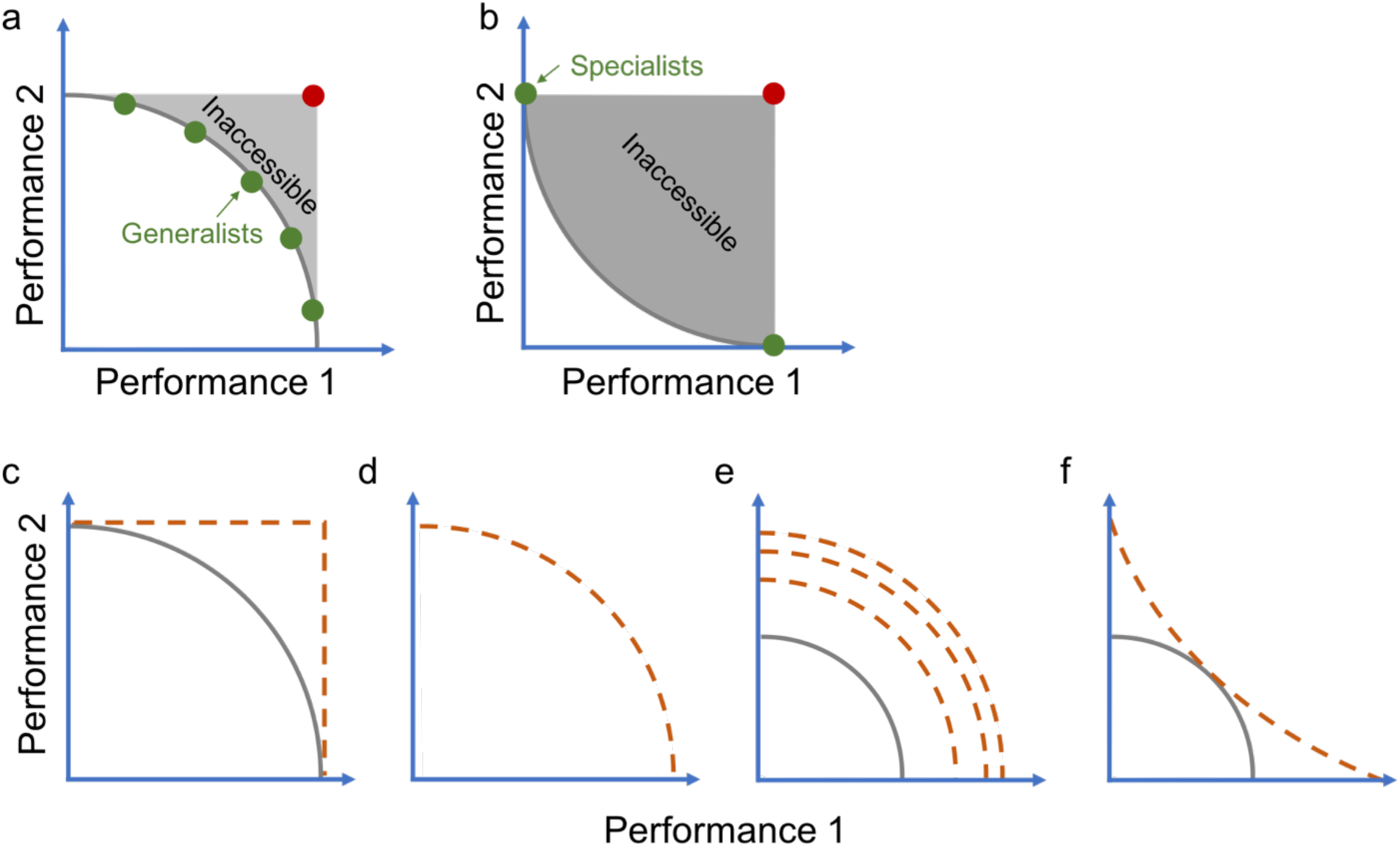
Pareto front geometry, and possible changes over longer-term evolution. **a-b,**(**a**) the convex-shaped Pareto front favors generalists, while (**b**) the concave-shaped front favors specialists during evolution. **c-f,** The current convex Pareto front (the solid grey curve) can (**c**) change into a rectangle, with the previously inaccessible space being populated, (**d**) stay in place, (**e**) move forward while keeping its shape, and (**f**) change its shape over longer-term evolution. Potential Pareto fronts after longer-term evolution are depicted in orange dashed lines.

Previous studies have used, for example, ecological data in phytoplankton^29^, interactions between phage and *E. coli*^30^, and synthetic, *E. coli* based systems^31^ to investigate the geometry of Pareto fronts, and in one case, it has been shown that an evolving ancestor is likely on a Pareto front^12^. However, no study has yet quantitatively defined a Pareto front or characterized its geometry in evolving populations where the ancestor lies behind the front, which is the case in most experimental evolutions. Here we identified not one, but two convex-shaped fronts for two independent tradeoffs under well-controlled selection pressures in our short-term evolution experiments. It is possible that the shape of the Pareto front itself may change over the timescales of evolution^32,33^ and the way in which it might change will be informative about whether the observed front is due solely to a genetic constraint, or instead whether there is an underlying intrinsic physiological constraint.

Over longer-term evolution, the space that is inaccessible in the short term may become populated, and the shape change to become a rectangle (Fig. 4c). This would imply there is no physiological constraint between the two traits and the observed Pareto front is purely due to a genetic constraint – that is, no clones with single mutations are able to occupy the seemingly inaccessible space, yet clones with multiple mutations can. Alternatively, the front may either stay in place (Fig. 4d), or move forward but retain the same shape (Fig. 4e), always defining an inaccessible space. This scenario would suggest intrinsic physiological constraints that no single individual could maximize performances in both traits simultaneously. A final possibility is that longer-term evolution may change the shape of the front from being convex to being concave (Fig. 4f) such that individuals with extreme performance in one or the other trait are the most fit depending upon the exact condition in which they are evolved.

The behavior of clones containing multiple adaptive mutations should provide some insights. We observed three clones carrying two adaptive mutations each in genes specific to different evolutionary conditions. These clones harbor mutations in *SXM1* and *HOG1, SXM1* and *SSK1*, and *SXM1* and *CYR1*, respectively. We observed that each of these double mutants is no closer to the front than the corresponding single mutants (Fig. S4), suggesting the front itself might be moderately stable. However, clearly both long-term evolution and further evolution of already adaptive clones under various conditions are needed to test this.

### Future Prospects

Despite much focus on the study of tradeoffs in ecology and evolution, rigorous demonstration of tradeoffs has proven surprisingly difficult^15,34^. Furthermore, even when tradeoffs have been demonstrated, the underlying causes typically remain elusive -- the genetic bases of adaptation and tradeoffs identified here provide additional potential targets for further investigation of whether the detected tradeoffs are caused by intrinsic physiological constraints. Here we have shown that it is possible to use barcoding and experimental evolution across a range of conditions to isolate a large enough number of adaptive mutants that together can map the shape of the evolutionary accessible trait space in short-term evolution, from which tradeoffs can be inferred. Our approach is generic and can be used to study tradeoffs between multiple traits including ecologically relevant traits such as the ability to sporulate or undergo mating and can be performed with different founding strains and species. Such studies hold promise in helping us to understand the shape of tradeoffs among multiple traits both in pairs and in higher dimensions.

## Data Availability

All sequencing data are deposited in Short Read Archive under Bioproject ID PRJNA515761.

## Acknowledgements

We wish to thank A. Agarwala, G. Kinsler, C. McFarland, and D. Fisher for discussion. We thank J. Blundell, K. Geiler-Samerotte, O. Kolodny, S. Kryazhimskiy, C. Li, F. Rosenzweig and S. Venkataram for comments on the manuscript. We thank all members in the Sherlock and Petrov labs for helpful suggestions. Y.L. is supported by the Stanford Center for Computational, Human and Evolutionary Genomics (CEHG) Predoctoral Fellowship. The work was supported by NIH grant R01GM110275 and NASA grant NNX17AG79G to G.S. and NIH grant R35GM118165 to D.A.P..

## Author Information

### Contributions

Conceptualization: Y.L., G.S., and D.A.P.. Methodology: Y.L.. Formal Analysis: Y.L.. Investigation: Y.L.. Writing – Original Draft: Y.L.. Writing – Review & Editing: Y.L., D.A.P., and G.S.. Supervision: D.A.P., and G.S..

## Supplementary Figures

**Figure S1:**
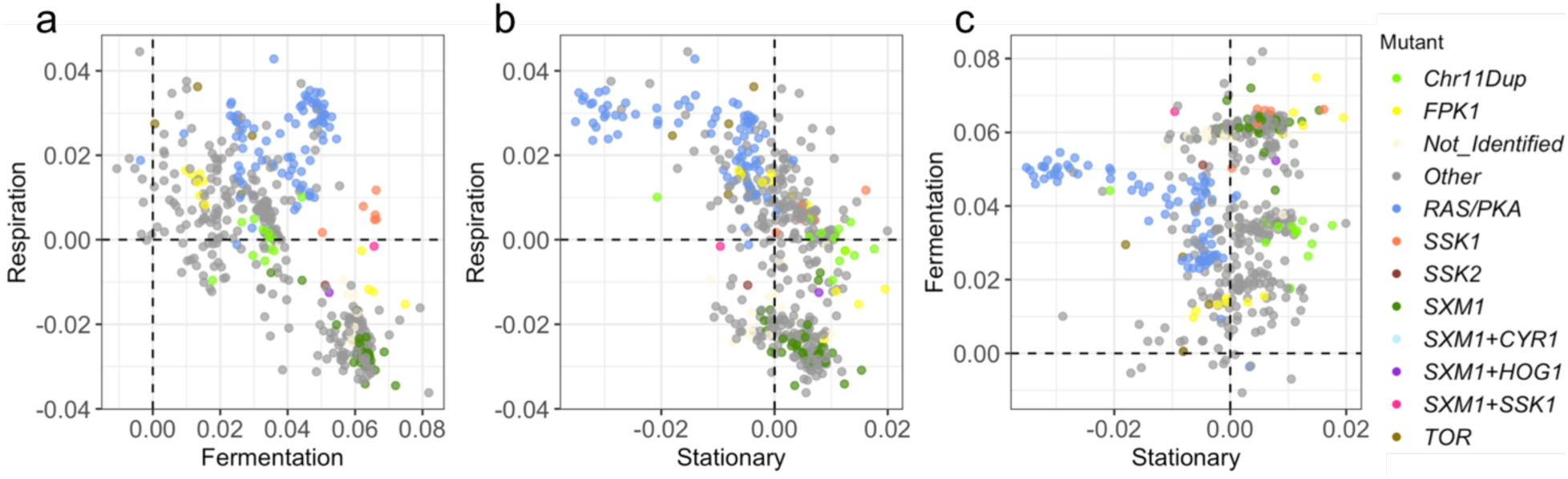
The genetic basis of adaptive clones in the trait space. **a-c,** Adaptive clones are colored by their genetic basis and plotted for each pair of performances. Each dot represents a lineage.

**Figure S2:**
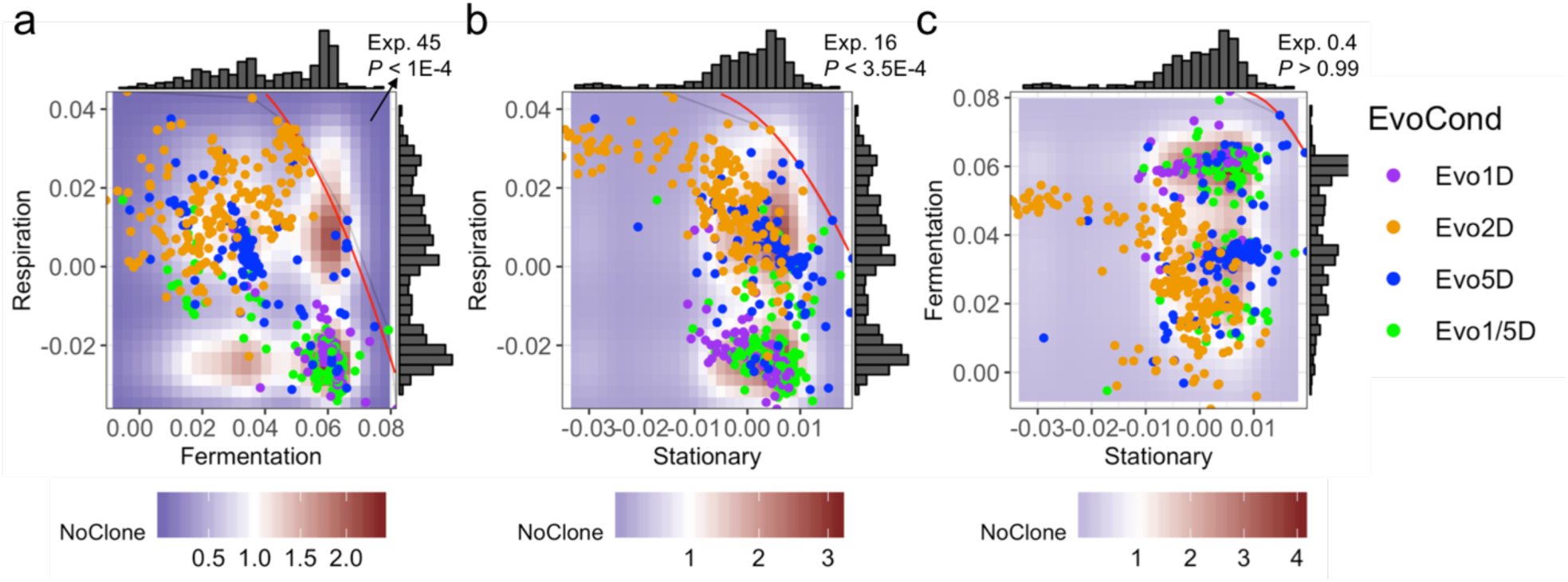
Null distribution indicates the existence of an evolutionarily inaccessible space. **a-c,** The background color represents the expected number of clones with corresponding performances under a null hypothesis that performances in different growth phases are independent. Clones, represented by dots, are colored by their evolutionary condition. The thin grey curves represent putative Pareto fronts drawn by the convex hull algorithm. The red curves represent the second degree polynomial fit of these putative Pareto fronts.

**Figure S3:**
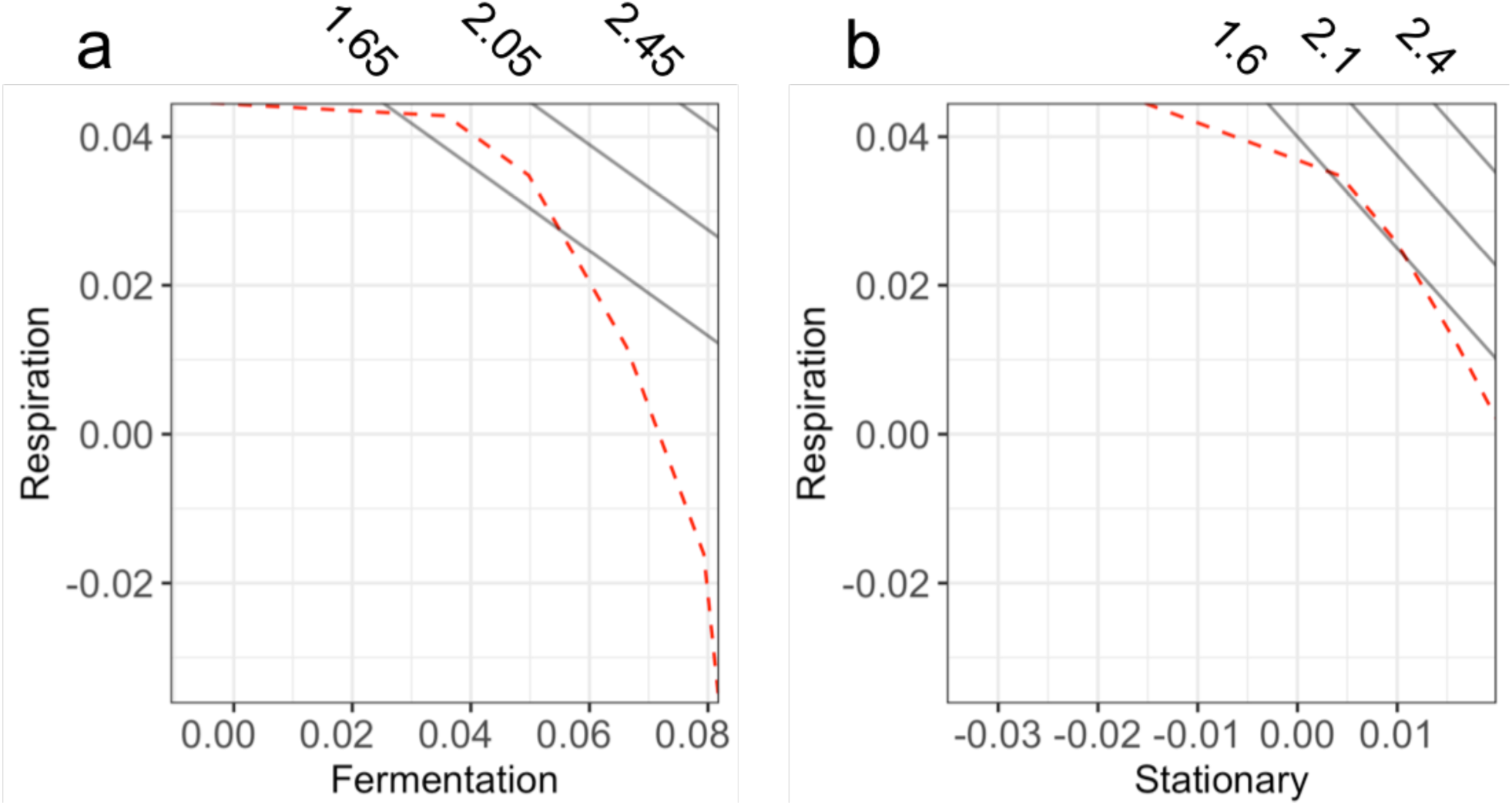
Fitness estimates in evolutionarily inaccessible space. **a-b,** The red dashed lines represent putative evolutionary fronts identified by the convex hull algorithm. The black lines represent (**a**) estimated fitness in Evo2D using the corresponding fermentation and respiration performance, and (**b**) estimated fitness in Evo5D using the corresponding respiration and stationary phase performance with the fermentation performance assumed to be zero. Fitness estimates per cycle are labeled on top of the panel.

**Figure S4:**
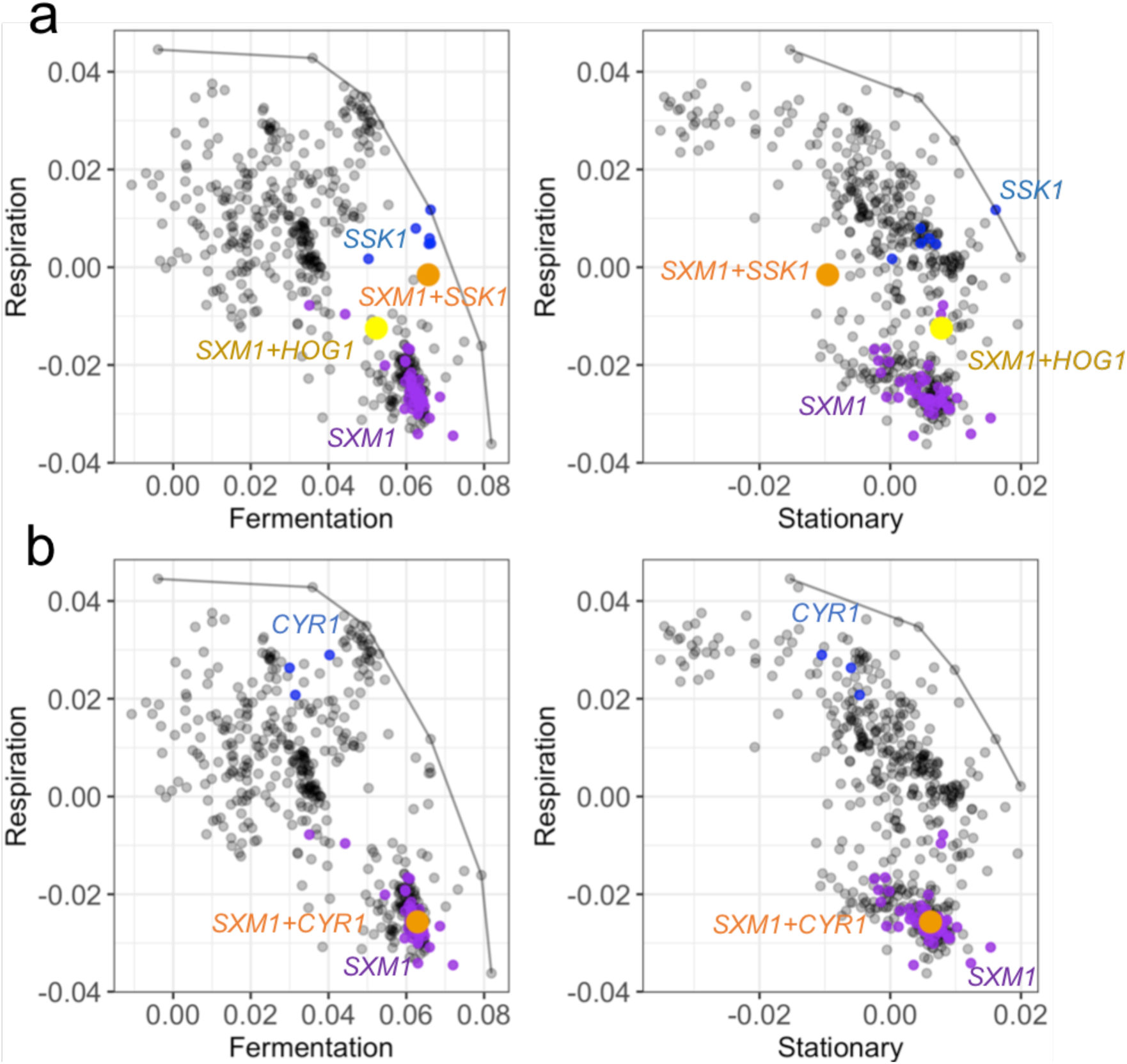
Epistasis between recurrent beneficial mutations. (**a**) Two mutants carrying mutations in both *SXM1* and HOG pathway genes, and (**b**) a mutant carrying mutations in both *SXM1* and RAS/PKA pathway gene *CYR1* are shown in the performance space. Double mutants are colored and shown in large dots. Their corresponding single mutants are colored and shown in small dots. Note that double mutants cannot outcompete both single mutants in all conditions and cannot break the detected Pareto fronts (in grey curve).

## Source Tables

Table S1: Barcode counts of all lineages during the course of evolution

Table S2: Fitness measurements of isolated clones

Table S3: Genetic basis of genome-wide sequenced clones

Table S4: Viability measurement of *FPK1* mutants and wild-type strains

## References

1. Loschiavo, S. R. Effect of oviposition sites on egg production and longevity of *Trogoderma parabile* (Coleoptera: Dermestidae). Can. Entomol. 100, 86–89 (1968).

2. Tinkle, D. W. The concept of reproductive effort and its relation to the evolution of life histories of lizards. Am. Nat. 103, 501–516 (1969).

3. Reznick, D. A., Bryga, H. & Endler, J. A. Experimentally induced life-history evolution in a natural population. Nature 346, 357–359 (1990).

4. The Evolution of Life Histories. (Oxford University Press, 1992).

5. Camargo, A., Sarroca, M. & Maneyro, R. Reproductive effort and the egg number vs. size trade-off in Physalaemus frogs (Anura: Leiuperidae). Acta Oecologica 34, 163–171 (2008).

6. Cunningham, J. T. Degenerative mutations. Nature 130, 203–204 (1932).

7. Wang, Y. et al. Contribution of both positive selection and relaxation of selective constraints to degeneration of flyability during geese domestication. PLOS ONE 12, e0185328 (2017).

8. Darwin, C. The descent of man, and selection in relation to sex. *Princeton University Press* (1871).

9. Shoval, O. et al. Evolutionary trade-offs, pareto optimality, and the geometry of phenotype space. Science 336, 1157–1160 (2012).

10. Tendler, A., Mayo, A. & Alon, U. Evolutionary tradeoffs, Pareto optimality and the morphology of ammonite shells. BMC Syst. Biol. 9, 12 (2015).

11. Mooney, K. A., Halitschke, R., Kessler, A. & Agrawal, A. A. Evolutionary trade-offs in plants mediate the strength of trophic cascades. Science 327, 1642–1644 (2010).

12. Fraebel, D. T. et al. Environment determines evolutionary trajectory in a constrained phenotypic space. eLife 6, e24669 (2017).

13. Nidelet, T. & Kaltz, O. Direct and correlated responses to selection in a host-parasite system: testing for the emergence of genotype specificity. Evol. Int. J. Org. Evol. 61, 1803–1811 (2007).

14. Buckling, A., Brockhurst, M. A., Travisano, M. & Rainey, P. B. Experimental adaptation to high and low quality environments under different scales of temporal variation. J. Evol. Biol. 20, 296–300 (2007).

15. Bono, L. M., Smith, L. B., Pfennig D. W. & Burch, C. L. The emergence of performance trade-offs during local adaptation: insights from experimental evolution. Mol. Ecol. 26, 1720– 1733 (2017).

16. McGee, L. W. et al. Synergistic pleiotropy overrides the costs of complexity in viral adaptation. Genetics 202, 285–295 (2016).

17. Jasmin, J.-N. & Kassen, R. On the experimental evolution of specialization and diversity in heterogeneous environments. Ecol. Lett. 10, 272–281 (2007).

18. Bennett A. F. & Lenski, R. E. An experimental test of evolutionary trade-offs during temperature adaptation. Proc. Natl. Acad. Sci. 104, 8649–8654 (2007).

19. Satterwhite R. S. & Cooper, T. F. Constraints on adaptation of Escherichia coli to mixed- resource environments increase over time. Evolution 69, 2067–2078 (2015).

20. Levy, S. F. et al. Quantitative evolutionary dynamics using high-resolution lineage tracking. Nature 519, 181–186 (2015).

21. Venkataram, S. et al. Development of a comprehensive genotype-to-fitness map of adaptation-driving mutations in yeast. Cell 166, 1585–1596.e22 (2016).

22. Li, Y. et al. Hidden complexity of yeast adaptation under simple evolutionary conditions. Curr. Biol. 28, 515–525.e6 (2018).

23. Seedorf, M. & Silver, P. A. Importin/karyopherin protein family members required for mRNA export from the?nucleus. Proc. Natl. Acad. Sci. U. S. A. 94, 8590–8595 (1997).

24. Yona, A. H. et al. Chromosomal duplication is a transient evolutionary solution to stress. Proc. Natl. Acad. Sci. 109, 21010–21015 (2012).

25. Natesuntorn, W. et al. Genome-wide construction of a series of designed segmental aneuploids in *Saccharomyces cerevisiae*. Sci. Rep. 5, 12510 (2015).

26. Sunshine, A. B. et al. The fitness consequences of aneuploidy are driven by condition- dependent gene effects. PLOS Biol. 13, e1002155 (2015).

27. Garay, E. et al. High-resolution profiling of stationary-phase survival reveals yeast longevity factors and their genetic interactions. PLOS Genet. 10, e1004168 (2014).

28. Levins, R. Theory of fitness in a heterogeneous environment. I. The fitness set and adaptive function. Am. Nat. 96, 361–373 (1962).

29. Ehrlich, E., Kath, N. J. & Gaedke, U. The shape of a defense-growth trade-off governs seasonal trait dynamics in natural phytoplankton. bioRxiv 462622 (2018). doi:10.1101/462622

30. Jessup C. M. & Bohannan, B. J. M. The shape of an ecological trade-off varies with environment. Ecol. Lett. 11, 947–959 (2008).

31. Maharjan, R. et al. The form of a trade-off determines the response to competition. Ecol. Lett. 16, 1267–1276 (2013).

32. Roff, D. A. & Fairbairn, D. J. The evolution of trade-offs: where are we? J. Evol. Biol. 20, 433–447 (2007).

33. Yi, X. & Dean, A. M. Phenotypic plasticity as an adaptation to a functional trade-off. eLife 5, e19307 (2016).

34. Sexton, J. P., Montiel, J., Shay, J. E., Stephens M. R. & Slatyer, R. A. Evolution of ecological niche breadth. Annu. Rev. Ecol. Evol. Syst. 48, 183–206 (2017).

