## Supplementary Materials and Methods for "Single Nucleotide Mapping of the Locally Accessible Trait Space in Yeast Reveals Pareto Fronts that Constrain Initial Adaptation"

### Supplementary Information

#### 1 Founding Populations and Experimental Evolution

##### 1.1 Barcoded yeast populations

Barcoded yeast populations were constructed as in Levy, Blundell *et al*[1]. However, where Levy, Blundell *et al*[1] used only one landing pad yeast strain (SHA185), we used multiple landing pad yeast strains, with each landing pad strain carrying a unique, condition-specific barcode. Next, a high-complexity plasmid library was introduced into each landing pad strain through transformation; correct integrants were selected for uracil prototrophy[1]. These transformants contain both a low-complexity *condition-specific* barcode and a high-complexity *lineage tracking* barcode. Three barcoded yeast populations with distinct condition-specific barcodes were constructed. Each barcoded yeast population includes around half a million unique transformants. These three barcoded yeast population were evolved under Evo1D, Evo5D and Evo1/5D, respectively.

##### 1.2 Yeast growth media and growth cycle

All cultures were grown in 100 mL of M3 medium[1] (5x Delft medium with 4% ammonium sulfate and 1.5% dextrose) in 500 mL Delong flasks (Bellco) at 30°C and 223 RPM. This culture condition is referred to as our standard culture condition.

Yeast cells go through different growth phases when growing under a glucose-limited condition, including the lag phase, fermentation, respiration and the stationary phase. Based on our previous study[2], with 5E7 cells inoculated into our standard culture condition, cells experience lag phase for ~4 hours, fermentation for ~16 hours, respiration for ~40 hours and stationary phase thereafter[2]. We note that these parameters can change somewhat under different growth conditions. For example, cells experience a longer lag phase under Evo5D since cells entered into stationary phase in the previous growth cycle. Such variations are not taken into account in our analyses.

##### 1.3 Serial-batch transfers during evolution

The barcoded yeast populations were evolved by serial batch culture in M3 medium. Under Evo1D, bottlenecks were performed by adding 7E7 cells of the culture from the previous growth cycle to fresh media; cells were grown for 1 day/24 hours between each bottleneck. Cells were highly viable (close to 100%) after 1 day of growth. Cell counts were performed at each bottleneck to estimate the generation time. Cells went through ~7 generations during each transfer in Evo1D. Populations were evolved for up to 25 growth cycles (~175 generations).

Under Evo5D, bottlenecks were performed every 5 day/120 hours. Cells' viability decreased during the stationary phase. To avoid strong bottleneck drift, the cells' viability was estimated at each bottleneck and 7E7 viable cells were inoculated to the next serial transfer. Viability was estimated by counting colony forming units (CFUs) on YPD plates and then dividing CFUs by the number of cells plated. Cells went through ~7.5 generations during each transfer in Evo5D. Cells were evolved up to 16 cycles (~120 generations).

Under Evo1/5D, cells were bottlenecked every 1 day and 5 days, alternating. 7E7 viable cells were inoculated at each bottleneck. Similarly, cells went through ~7 generations during 1-day transfer and ~7.5 generations during 5-day transfer. Cells were evolved up to 23 cycles by alternating between 1 day and 5 day transfers (~166 generations).

Two replicates were conducted under each evolutionary condition. At the end of each transfer, 2mL cell culture of evolution were mixed with 1ml 40% glycerol, aliquoted into two Eppendorf tubes and stored at -80°C. The rest of the cell culture (>95mL, besides a small amount used to inoculate the next cycle), was centrifuged, then the cell pellet was resuspended in 5mL sorbitol solution (0.9M sorbitol, 0.1M Tris-HCL pH 7.5, 0.1M EDTA pH 8.0), and then aliquoted into 2mL Eppendorf tubes and stored at -20°C for genomic extraction.

#### 2 Tracking Lineage Dynamics During Evolution

##### 2.1 Library preparation and sequencing

To track lineages' frequency changes over the course of evolution, cells collected at the end of every three transfers under Evo1D, Evo5D and Evo1/5D were used for genomic DNA extraction. Genome extraction and PCR amplification of the barcode region were conducted as in Venkataram, Dunn *et al*[3]. Amplicons were sequenced on Illumina HiSeq 4000 (2X101 paired end). Data for Evo2D can be found in Levy, Blundell et al[1]. A perfect sequencing read should follow this DNA sequence:

NNNNNNNNXXXXXTTAATATGGACTAAAGGAGGCTTTTGTCGACGGATCCGATATCGGTACC (+26bp lineage Barcode+) ATAACTTCGTATAATGTATGCTATACGAAGTTAT (+26bp condition barcode+)

GGTACCGATATCAGATCTAAGCTTGAATTCGATXXXXXXXXXNNNNNNNN

The Ns in the sequence are random nucleotides and are used as UMIs in the downstream analysis to remove skew in the counts caused by PCR jack-potting[1]. The Xs correspond to multiplexing tags, which allow different samples to be distinguished when loaded on the same sequencing flow cell[1].

##### 2.2 Barcode counts

Sequencing reads, with 17bp (including the UMIs and Multiplexing tags) on both the 5' and the 3' end removed, were first aligned to the reference barcode region using Bowtie2. Based on the alignment, both condition and lineage barcodes were extracted. Next, the UMIs, Multiplexing tags and barcodes were re-associated for each pair of reads. Reads were split into different files based on their Multiplexing tags, representing which sample the reads came from. Lastly, reads from the same sample were clustered using Bartender[4] with Hamming distance 2 and seed length 8. The final output includes both barcode sequences and counts of each lineage. Each lineage is represented by a unique combination of condition and lineage barcodes. Reads with the same combination of UMI, Multiplexing tags and barcodes were counted as one read as they are likely caused by PCR jack-potting. An extra round of clustering was conducted using barcode combinations identified by Bartender to further cluster barcodes with <=2 Hamming distance. On average, each sample has ~1.3 million high-quality reads. For each evolution replicate, lineages with fewer than 10 counts across all time points and lineages present at fewer than three sequenced time points were filtered out. Barcode counts over the course of evolution can be found in Table S1.

#### 3 Isolated Clones from Evolution

##### 3.1 Isolate evolved clones from evolutionary conditions

Based on population dynamics, yeast clones were isolated from frozen samples of Evo1D, Evo5D and Evo1/5D at cycle 11. ~50μL of frozen stock (containing ~3.5E6 cells) was diluted into 500μl PBS plus 1μl propidium iodide and used for flow cytometry sorting. Single cells were sorted into 96-well plates with 100μL YPD media in each well. 16 plates of cells (~1,500 cells) were sorted for each evolutionary condition with 8 plates from each replicate (48 plates in total sorted from Evo1D, Evo5D and Evo1/5D). Sorted cells were grown at 30°C for three days without shaking and reached saturation by day three. ~5μL of saturated cell culture were removed from each well and inoculated into a different 96-well plate with 100μL fresh YPD media in each well. These replicated plates were grown at 30°C for two days without shaking to reach saturation and used for following ploidy and barcode identification. The rest of the saturated cell culture was mixed with 100μL 50% glycerol and stored at -80°C.

**3.2 The ploidy test**

A high throughput ploidy test was developed using the drug benomyl[3]. The saturated cultures from above 48 replicated 96-well plates were mixed and pinned onto YPD+20μg/ml benomyl (in DMSO) rectangular agar plates using a multi-pronged pinner, grown at 25°C for 48 hours, and then imaged. Under these conditions, diploid growth is strongly inhibited by benomyl but haploid growth is less affected.

**3.3 DNA barcode identification by Metagrid**

20μL of saturated cell culture were removed from each well of the replicated plates into 96-well PCR plates. Cells were lysed by incubating at 95°C for 15 mins. 5μL of lysed cell culture were used to PCR amplify the DNA barcode region. To reduce the cost of Sanger sequencing ~4,600 clones, a Metagrid approach was developed. 72 forward and 64 reverse primers were synthesized with each primer carrying an 8bp unique multiplexing tag (Ns in the primer sequence). Each PCR well was coded by a unique combination of forward and reverse multiplexing tags. Two-step PCR amplification was conducted here with the first step using multiplexed primers and the second step using standard Illumina paired-end ligation primers (PE1 and PE2). PCR products from all 48 plates were pooled together and sequenced using Illumina NextSeq (2X150 paired end).

Forward Primer of the First Step PCR:

ACACTCTTTCCCTACACGACGCTCTTCCGATCTNNNNNNNNTTAATATGGACTAAAGGAGG CTTTT

Reverse Primer of the First Step PCR:

CTCGGCATTCCTGCTGAACCGCTCTTCCGATCTNNNNNNNNTCGAATTCAAGCTTAGATCT GATA

Regular expressions that match DNA sequences flanking multiplexing tags of primers and clones' barcodes were used to extract multiplexing tags and barcodes. A map between clones' DNA barcodes and their physical positions on the 96-well plates was constructed based on multiplexing tags. The number of clones carrying unique barcodes is lower than the number of clones isolated due to the lack of growth in a small number of wells and more importantly, the high frequency of a few adaptive clones -- a large number of isolated clones carrying the same barcodes. 124, 166, and 397 unique barcodes were identified from Evo1D, Evo5D and Evo1/5D respectively.

#### 4. High-throughput Fitness Measurements

##### 4.1 Pool of clones for fitness measurements

Isolated clones with unique barcodes were pooled together for high-throughput fitness assays. Note that some barcodes were present in both haploid and diploid clones; for these barcodes, only haploid clones were pooled for fitness assays. Clones with unique barcodes were hand-picked from previous frozen stock and re-arrayed onto a set of 96 deep-well plates with 700μL YPD medium in each well. Cells were grown at 30°C for 2 days to reach saturation without shaking. 500μL of 50% glycerol were added into each well using a multichannel pipette. 1mL of the mixture from each well was pooled together into a container. The pooled cell culture was mixed well and aliquoted into 2mL Eppendorf tubes, which were stored at 80°C for future fitness measurements.

##### 4.2 Fitness measurement conditions

We measured clones' fitness under four conditions, 1-day, 2-day, 3-day and 5-day serial batch culture conditions (Fit1D, Fit2D, Fit3D and Fit5D). These four conditions are the same except the length of the growth cycle. Out of which, 1-day, 2-day and 5-day serial batch culture conditions are the same as Evo1D, Evo2D and Evo5D evolutionary conditions. Note that fitness of Evo2D clones were measured in Li, Venkataram *et al*[2]. Clones isolated from Evo1D, Evo5D, and Evo1/5D were measured in this study and analyzed in this section. Note that all data reported in this section can be found in Table S2.

##### 4.3 Preculture

A tube of pooled cell culture was removed from -80°C and thawed at room temperature. 1mL of pooled cell culture was inoculated into 15ml fresh M3 media contained in a 500ml Delong flask and grown at 30°C shaking at 223 RPM overnight for cell propagation. 400µL of overnight cell culture were inoculated into 100mL fresh M3 media (X4) and precultured at the standard condition.

An ancestral clone carrying a restriction site in the barcode region was previously constructed and used to compete with evolved clones for fitness measurements[3]. The ancestor clone was streaked out from a freezer stock onto M3 agar plates and grown for 2 days until colonies were visible. A single colony was inoculated into 3mL of M3 medium and grown for 48 hours (30°C roller drum). After saturation, 400µL were used to inoculate pre-cultures (100mL M3 media in 500mL Delong flasks, 223 RPM 30°C). Like the pooled cell culture, four pre-cultures were prepared.

The pooled preculture and the ancestor preculture were acclimated for the same cycle length as that of the fitness measurement condition. For instance, precultures were grown for one day before being mixed and assayed under the 1-day fitness measurement condition.

##### 4.4 High-throughput competition

Fitness assays were conducted by mixing the pooled preculture with the ancestor preculture in a 1:9 ratio (time 0) and growing this mixed culture for four successive growth cycles (time points 1, 2, 3 and 4). In Fit1D, at the end of each cycle, 450ul cell culture was inoculated into 100ml fresh media to start the next cycle. In Fit2D, Fit3D and Fit5D, at the end of each cycle, 400ul cell culture was inoculated. Cells were collected at time 0, and at the end of each of the four growth cycles. Cell pellet from each sample was resuspend in 5ml sorbitol solution (0.9M sorbitol, 0.1M

Tris-HCL pH 7.5, 0.1M EDTA pH 8.0), aliquoted into 2mL Eppendorf tubes and stored at -20°C. Three replicates were performed under each fitness assay condition. Genome extraction, barcode region amplification and Illumina sequencing were conducted for each sample.

**4.5 Library preparation of high-throughput fitness assays**

Genomic DNA was extracted with following steps: 1) remove collected cells from -20°C and thaw at room temperature; 2) spin down cells and wash them with water; 3) resuspend cells in 400µL buffer (0.9 M Sorbitol, 50mM Na Phosphate pH 7.5, 240µg/mL zymolase, 14mM ß-mercaptoethanol) and incubate at 37°C for 30 minutes; 4) add 40µL 0.5M EDTA and vortex; 5) Add 40µL 10% SDS and vortex; 6) add 56µL 20mg/mL proteinase K (Life Technologies 25530-015), vortex very briefly and incubate at 65°C for 30 minutes; 7) incubate samples on ice for 5 minutes; 8) add 200µL 5M potassium acetate, shake and incubate on ice for 30 minutes; 9) spin for 10 minutes, transfer supernatant to a new tube with 750µL ispropanol and rest on ice for 5 minutes; 10) spin down for 10 minutes and wash twice with 70% ethanol; 11) resuspend in 50µL 10mM Tris pH 7.5 -- leave on bench overnight if pellet is not resuspended fully; and 12) add 0.5µL 20 mg/mL RNase A and incubate at 65°C for 30 minutes.

Two-step PCR was used to amplify the barcode region. See Venkataram, Dunn *et al*[3] for primer details. The first step PCR was conducted using 6µg genomic DNA and separated into six 50µL reactions.

| | |
|---|---|
| OneTaq 2X Mix | 150µL |
| Forward Primer 5uM | 3µL |
| Reverse Primer 5uM | 3µL |
| Template | 6µg |
| MgCl2 50mM | 12µL |
| H20 | add up to 300µL |

PCR program for the first step PCR:

| | | |
|---|---|---|
| 1X | 94°C | 10 minutes |
| 3X | 94°C | 3 minutes |
| | 55°C | 1 minute |
| | 68°C | 1 minute |
| 1X | 68°C | 1 minute |

| | | |
|---|---|---|
| | 4°C | inf |

We then performed PCR cleanups following the QIAquick PCR purification protocol (6 PCR reactions pooled into one column) and elute into 30μL $H_2O$.

The second step PCR was completed in one reaction using Herculase II fusion DNA polymerase (Agilent 600677):

| | |
|---|---|
| Buffer | 10μL |
| dNTP 100μM | 0.5μL |
| Herculase | 0.5μL |
| PE1 10μM | 1.25μL |
| PE2 10μM | 1.25μL |
| Template | 30μL |
| $H_2O$ | 6.5μL |

PCR program for the second step PCR:

| | | |
|---|---|---|
| 1x | 98°C | 2 minutes |
| 20x | 98°C | 10 seconds |
| | 69°C | 20 seconds |
| | 72°C | 30 seconds |
| 1x | 72°C | 1 minute |
| | 4°C | inf |

**4.6 Fitness estimates**

The number of DNA barcodes was tracked by Illumina sequencing (Illumina NextSeq), which was then used to estimate lineages' frequencies in the population as a whole, as previously described[2,3]. In this study, times 0, 1, 2, and 3 were used for fitness estimates under Fit1D, Fit2D and Fit3D. Time 0, 1, 2, 3, and 4 were used for fitness estimates under Fit5D. The source code for computing these fitness estimates can be found at https://github.com/barcoding-bfa/fitness-assay-python. First, we input barcode counts to the script. After the 1$^{st}$ run, the script output barcodes that were likely to be neutral. During the 2$^{nd}$ run, we input both the barcode counts and a list of neutral barcodes estimated from the 1$^{st}$ run. Fitness estimates from the 2$^{nd}$ run were used for further analysis. Final fitness estimates were calculated by inverse variance weighting of estimates from all three replicates. Barcodes identified in these fitness assays were mapped back to barcodes identified in Metagrid. Thus, for each unique barcode, we know the fitness values under all fitness measurement conditions, its physical position in a 96-well plate in frozen stock, and the associated ploidy. Note that the physical position of each barcode is required for picking clones for genome-wide sequencing. In sum, 661 unique barcodes had confidently called ploidy and were successfully identified from our fitness assays. Out of the 661 unique barcodes, 644 of them had high-quality fitness values under every fitness assay condition and were used for further analysis.

#### 5 Classification of Clones

The 644 lineages with high-quality fitness measurements were classified into four groups, based on their ploidy and fitness: neutral haploids, adaptive haploids, diploids presumed to have no additional adaptive mutations (“pure” diploids), and diploids with additional adaptive mutations (high fitness diploids; see Li, Venkataram *et al*[2] for details). Briefly, adaptive haploids were defined as lineages that had low probability ($p < 1e-3$) of having log fitness of 0 or less in at least two conditions; high fitness diploids were defined as diploid lineages with additional adaptive mutations that had low probability ($p < 1e-3$) of having fitness less than the mean fitness of the diploid class in at least one condition. The mean fitness of the diploid population was calculated using inverse variance weighting. In total, 254 adaptive haploids, 66 high fitness diploids, 218 neutral haploids and 106 pure diploids were characterized among clones isolated from Evo1D, Evo5D, and Evo1/5D. See Li, Venkataram *et al*[2] for classification of Evo2D clones.

#### 6 Quantification of Performances in Growth Phases

Fitness change per hour was used as the measurement of lineages’ performances in different growth phases. Respiration performance per hour was calculated by subtracting Fit1D measurements from Fit2D measurements and dividing by 24 hours (because compared to Fit1D, clones experienced 24 hours of extra respiration in Fit2D, while keeping the lag and fermentation phases roughly the same). Fit1D almost directly measures the performance in fermentation but did contain ~4h respiration per transfer. Therefore, using above estimates of respiration performance per hour, we subtracted expected fitness change in the extra 4 hours of respiration from Fit1D measurements. We then divided the difference by 16 hours of fermentation to estimate fermentation performance per hour. Lastly, stationary phase performance per hour was calculated by subtracting Fit3D from Fit5D measurements and dividing the difference by 48 hours of stationary phase. The same quantification was performed for Evo2D clones, using their fitness measured under 1-day, 2-day, 3-day and 5-day serial transfer conditions[2]. See Li, Venkataram *et al*[2] for statistical details.

#### 7 Genome-wide Sequencing and Variant Calling

##### 7.1 Genome-wide sequencing library preparation

Clones selected for sequencing were grown in 500µL YPD on 96 deep-well plates for two days at 30°C without shaking. 400µL of cell culture were collected from each well for DNA extraction. Genomic DNA was prepared using Invitrogen PureLink Pro 96 Genomic DNA Kit (Catalog no. K1821-04A) in a 96-well format. Libraries were constructed and multiplexed using Nextera technology with the protocol of Kryazhimskiy, Rice *et al*[5]. Samples were sequenced with 2x150 Illumina Next-seq paired end sequencing technology. 179 adaptive haploids, 20 high fitness diploids, 8 neutral haploids and 9 pure diploids were sequenced with an average coverage >50 for both haploids and diploids. Note that only clones isolated from Evo1D, Evo5D, and Evo1/5D are sequenced and analyzed in this study and reported in this section. See Venkataram, Dunn *et al*[3] and Li, Venkataram *et al*[2] for details about Evo2D clones.

##### 7.2 FASTQ processing

For each sample, we received two fastq files, one for each read of the paired end sequencing (“fastqR1” and “fastqR2”). Using cutadapt 1.16, we trimmed the first 10bp of each read (-u 10), low-quality ends (-q 30) and any adapter sequences (-a). After trimming, sequences with a length shorter than 12bp (--minimum-length 12) were discarded.

First trim the forward read, writing output to temporary files (note, commands are a single line):

```
cutadapt --minimum-length 12 -q 30 -u 10 -a
CTGTCTCTTATACACATCTCCGAGCCCACGAGAC -o tmp.1.fastq.gz -p tmp.2.fastq.gz
fastqR1 fastqR2
```

Then trim the reverse read, using the temporary files as input:

```
cutadapt --minimum-length 12 -q 30 -u 10 -a
CTGTCTCTTATACACATCTGACGCTGCCGACGA -o trimmedR2.fastq.gz -p
trimmedR1.fastq.gz tmp.2.fastq.gz tmp.1.fastq.gz
```

Reads were mapped using bwa to *S. cerevisiae S288C* reference genome R64-1-1 (https://downloads.yeastgenome.org/sequence/S288C_reference/genome_releases/) and sorted using Sentieon Genomic Tools[6].

```
(bwa mem -M -R readGroupInfo -K 10000000 ReferenceGenome trimmedR1.fastq.gz trimmedR2.fastq.gz) | sentieon util sort -o SORTED_BAM --sam2bam -i –
```

Duplicates were removed using the sorted BAM file. The first command collected read information, and the second command performed the deduping.

```
sentieon driver -i SORTED_BAM --algo LocusCollector --fun score_info SCORE_TXT
```

```
sentieon driver -i SORTED_BAM --algo Dedup -- rmdup --score_info SCORE_TXT --metrics DEDUP_METRIC_TXT DEDUP_BAM
```

Local realignment around indels was performed using the deduped BAM file.

```
sentieon driver -r ReferenceGenome -i DEDUP_BAM -- algo Realigner REALIGNED_BAM
```

Lastly, base quality score recalibration was performed using the realigned BAM file. The first command calculated the required modification of the quality scores assigned to individual read bases of the sequence read data. The second command applied the recalibration to calculate the post calibration data table.

```
sentieon driver -r ReferenceGenome -i REALIGNED_BAM --algo QualCal RECAL_DATA.TABLE
```

```
sentieon driver -r ReferenceGenome -i REALIGNED_BAM -q RECAL_DATA.TABLE --algo QualCal RECAL_DATA.TABLE.POST
```

##### 7.3 SNP and small Indel variant calling

SNP and small indels variants were called by the DNAscope algorithm (Sentieon Genomic Software) using the realigned BAM file and the output table of the base quality score recalibration (RECAL_DATA.TABLE). The parameter ploidy is assigned as 1 for haploids and as 2 for diploids.

```
sentieon driver -r ReferenceGenome -i REALIGNED_BAM -q RECAL_DATA.TABLE --algo DNAscope --ploidy [1|2] VARIANT_VCF
```

##### 7.4 Structural variant calling

The first command enabled the DNAscope algorithm to detect the break-end variant type (BND). The parameter ploidy was assigned as 1 for haploids and as 2 for diploids. The second command processed the temporary VCF file using the SVSolver algorithm and output structural variants to a VCF file.

```
sentieon driver -r ReferenceGenome -i REALIGNED_BAM -q RECAL_DATA.TABLE --algo DNAscope --var_type bnd --ploidy [1|2] TMP_VARIANT_VCF
```

Then process the VCF using the SVSolver algorithm with the following command:

```
sentieon driver -r ReferenceGenome --algo SVSolver -v TMP_VARIANT_VCF STRUCTURAL_VARIANT_VCF
```

##### 7.5 Copy number variant (CNV) calling

First, recalibrated BAM files were created using previously generated aligned BAM files and quality score calibration tables.

```
sentieon driver -r ReferenceGenome -i REALIGNED_BAM -q RECAL_DATA_TABLE --algo QualCal RECAL_DATA_TABLE_POST --algo ReadWriter RECALIBRATED_BAM
```

Second, the Panel of normal (OUT_PON) was created using recalibrated BAM files. By changing the window size of the bed file, the window size used to call CNVs was varied. Window sizes (200bp and 10kb) were applied in this study. The PON files were created only with the sequencing of haploids.

```
sentieon driver -r ReferenceGenome -i RECALED_BAM_1 [-i RECALED_BAM_n] --algo CNV --target BED_FILE --create_pon OUT_PON
```

Third, for each strain, the CNVs were called using the strain's recalibrated BAM file, the bed file and the PON file.

```
sentieon driver -r ReferenceGenome -i RECALED_BAM --algo CNV --target BED_FILE --pon PON_FILE OUT_CNV
```

##### 7.6 Variant annotation

Here the vcf file from SNP and small indel variants calling (VARIANT_VCF) is used as an example. Same commands were used to annotate structural variants.

Use snpEff[7] (http://snpeff.sourceforge.net/download.html) to annotate a vcf file and output the annotated vcf file, named Ann.vcf.

```
java -Xmx2g -jar snpEff -c snpEff_config -v R64-1-1.75 -class VARIANT_VCF > Ann.vcf -s snpEff_summary.html
```

For variants in coding regions, SNPSift was used to extract the first annotation of each variant, which is the annotation with the largest effect. Output the extracted annotation as a vcf file, named Final_Ann.vcf:

```
java -jar SnpSift extractFields Ann.vcf CHROM POS ID REF ALT QUAL FILTER EFF[0].EFFECT EFF[0].GENE: EFF[0].IMPACT: EFF[0].FUNCLASS: EFF[0].CODON: EFF[0].AA ANN[0].BIOTYPE: GEN[0].GT GEN[0].AD GEN[0].DP GEN[0].GQ GEN[0].PL > Final_Ann.vcf
```

For variants in non-coding regions, the nearest gene of each variant was extracted. Thus, the non-coding variants were annotated as either the upstream or downstream of the nearest genes.

##### 7.7 Filtering SNPs, small indels and structural variants

First, mitochondrial variants were discarded. Second, background variants, present in all strains, were removed. Variants present in >10 clones, which are isolated from more than one evolutionary condition, were also considered as background variants and discarded. Third, any variants in genes *FLO1* and *FLO9* were filtered out due to poor alignment in both genomic regions. Fourth, variants with a quality score smaller than 200 were filtered out. Note that if a variant was present in multiple clones, the alignment of this variant was manually checked regardless of its quality score and a decision was made based on all clones carrying this variant. Thus, a variant with a quality score <200 may not be filtered out if the same variant contained in other clones was proven to be authentic. Similarly, a variant with a quality score >200 may be filtered out if the same variant was proven to be bogus in other clones. Only 6 out of ~500 variants had quality scores <200. Lastly, by manually checking BAM files after alignment, variants within repetitive regions and regions with a poor alignment were filtered out. In addition, if a variant is present in multiple clones and these clones carry a same condition barcode, this variant is likely to be pre-existing mutations introduced during the barcoded population construction. These pre-existing mutations can be causative mutations and thus are kept in our analysis.

##### 7.8 Filtering copy number variants (CNVs)

Our DNA extraction and sequencing protocols resulted in an unevenly distributed and relatively low coverage of short chromosomes, including chromosomes I, III and VI. Thus, CNVs associated with these three chromosomes were not considered. For both haploids and diploids, CNVs generated with a 10kb window size were used to identify large duplications and deletions, combined with visual inspection of coverage plots. These variants were further confirmed by CNVs generated with a 200bp window size.

#### 8 Detailed Information of Mutations

179 adaptive haploids, 20 high fitness diploids, 8 neutral haploids and 9 pure diploids from Evo1D, Evo5D, and Evo1/5D were sequenced in this study, with 163 adaptive haploids, 19 high fitness diploids, 8 neutral haploids, and 7 pure diploids have variants identified. All detailed information of variants can be found in Table S3.

Pre-existing mutations were identified as 1) mutations in the same gene that are the exact same alleles, and 2) clones carrying identical mutations in the same gene with have the same condition barcodes. Thus, these pre-existing mutations were likely introduced during the construction of barcoded yeast populations. All 8 sequenced neutral haploids and 1 out of 9 sequenced pure diploids harbor chromosome IX duplication, suggesting that chromosome IX duplication is not an adaptative event. Since lineages with chromosome IX duplications have different condition barcodes, it is likely that chromosome IX duplication independently happened during the population construction. In this work, chromosome IX duplications are treated as pre-existing mutations. Note that even though the same pre-existing mutation appear in multiple clones, it is regarded as one independent adaptive event. The pre-existing mutations are not necessarily non-adaptive. For instance, despite pre-existing *SSK1* mutations, *SSK1* mutation also independently happened during evolution and thus is very likely to be adaptive. In addition, *LSM2, NUT2, TFB3, tL(GAG)G* and *VOA1* genes harbor both pre-existing mutations and mutations independently happened during evolution.

Lineages carrying mutations in *SSK1*, *SSK2* or *HOG1* gene are high-osmolarity glycerol (HOG) pathway mutants. Lineages carrying mutations in Ras/PKA pathway genes or TOR/Sch9 pathway genes are referred to as nutrient response pathway mutants in this work: *RAS2, GPB1, GPB2, PDE2, IRA1, CYR1, TFS1* and *YAK1* gene are involved in Ras/PKA signaling pathway; *SCH9, TOR1, KOG1* and *MDS3* are involved in TOR/Sch9 signaling pathway.

Recurrent mutations within genes or pathways are highly unlikely under neutral evolution and are a hallmark of adaptive mutations. Genes or pathways that were independently mutated five times or more were reported in Table 1 in the main text, including the duplication of chromosome XI that appeared more than five times. The full list of genes that were independently hit more than once can be found in Table S3. None of these multi-hit genes/pathways were mutated in neutral haploids and pure diploids sequenced in this study. By contrast, these multi-hit genes/pathways were mutated in 118 out of 182 adaptive clones with identified variants (~65%). In addition, 79 out of 182 adaptive clones (~43%) harbor mutations in genes/pathways that were independently hit more than five times.

Three lineages carry mutations in coding regions of two genes listed in Table 1 in the main text, offering an opportunity to study epistasis among these beneficial mutations. Specifically, these three lineages carry mutations in coding regions of *SXM1* and *SSK1*, *SXM1* and *HOG1* and *SXM1* and *CYR1*, respectively (Fig. S4). These three lineages are not colored in Fig. 3a-c in the main text. In addition, diploids with chromosome XI duplication and clones with *SSK2* mutations from Evo2D are not colored in Fig. 3a-c in the main text. If a mutant harbors more than one mutation, as long as one and only one of the mutations is located in genes listed in Table 1, it is classified as a mutant carrying mutations in genes listed in Table 1 (Fig. S1).

#### 9 Identification and Fitting of Pareto Optimality Fronts

The fermentation, respiration and stationary performances of adaptive haploids and high fitness diploids from Evo1D, Evo2D, Evo5D, and Evo1/5D were used for the following analysis. The algorithm “convex hull” was used to identify the smallest set of points (the convex set) that when connected, enclosed the rest of points. By connecting the two points with the largest value on x or y, a line was formed. Out of the convex set, points either on or above this line were used to depict the Pareto front. 6, 5 and 3 points were identified to depict the Pareto front between fermentation and respiration performance, between respiration and stationary phase performance and between fermentation and stationary phase performance, respectively.

With known points on Pareto fronts, regression was used, looking for a curve that fit the front. First to third degree polynomials were applied to the front between fermentation and respiration performance (adjusted r2: 0.64, 0.96 and 0.97 for 1$^{st}$, 2$^{nd}$ and 3$^{rd}$ degree polynomial fit, respectively) and between respiration and stationary phase performance (adjusted r2: 0.81, 1 and 1). In both cases, second degree polynomials offered a good fit and characterized curves were proven to be convex. Due to the small number of points on the front (including 3 points) between fermentation and stationary phase performance, we were unable to compare the first degree polynomial fit to the second degree polynomial fit.

#### 10 Null Distribution of Clones in the Inaccessible Space

We tested whether observed Pareto fronts and evolutionary inaccessible space were generated by random processes. For each pair of performances, the evolutionary inaccessible space was defined as the empty space between the Pareto front (the second polynomial fit) and the upper limits of both performances. The null hypothesis assumes that a pair of performances are independent from each other. We sampled the two tested performances independently with replacement for 504 times (the total number of adaptive clones from all four evolutionary conditions) and calculated the number of clones appearing in the inaccessible space under the null hypothesis. Using bootstrap, we repeated the test for 10,000 times and calculated the average number of clones expected to be observed in the inaccessible space (Mean) under the null hypothesis. Since the curve was not perfectly fit, a small number of clones was observed to be above or along the curve (<4 clones in fermentation and respiration space, <1 clone in respiration and stationary space, and 0 clones in fermentation and stationary space), referred to as the observed number of clones in the inaccessible space (Observed). A p-value was calculated as the probability of getting something equally or more extreme than what we observed under the null distribution. So the p-value was the probability of being more than abs(Mean-Observed) units away from the Mean when testing for 10,000 times, which is P(Test ClonesInAccessibleSpace <= Observed) + P(Test ClonesInAccessibleSpace >= Mean+abs(Mean-Observed)).

We then calculated the minimum number of clones that is needed to define the inaccessible space. By random sampling current adaptive clones, we concluded that at least 82 and 250 clones are required to confidently ($P < 0.01$) define the inaccessible space between fermentation and respiration performance, and between respiration and stationary phase performance, respectively.

#### 11 Mathematical Model to Calculate the Probability of not Sampling Optimal Types

##### 11.1 Equations

By knowing the selection coefficient (*s per generation*) of a particular adaptive event, the bottleneck population size ($N_b$), the mutation rate ($\mu$) and the effective variance of cell division per generation ($2c$), we can estimate the density distribution of establishment time ($\tau$) of this adaptive event (equation 7 in Supplementary Information of Levy, Blundell *et al*[1]). This density distribution represents for a given time point the probability that an adaptive event occurs and rises to $c/s$ cells to avoid drift (see Levy, Blundell *et al*[1] for details).

$$p(\tau)\, d\tau = s/\Gamma(N_b{*}\mu/c) * Exp[ -N_b{*}\mu{*}(s/c){*}\, \tau - Exp(-s{*}\tau) ]$$

After establishment, clones grow exponentially (cell number $n = c/s{*}Exp[s{*}(t-\tau)]$ with a given establishment time $\tau$ at a sampled time $t$). Thus, the frequency of the adaptive event at a sampled time ($t$) can be estimated using the density distribution of establishment time.

$$f = \int_0^t d\tau * s/\Gamma(N_b{*}\mu/c) * Exp[ -N_b{*}\mu{*}(s/c){*}\, \tau - Exp(-s{*}\tau) ] * c/(N_b{*}s) * Exp[s{*}(t-\tau)] ]$$

Lastly, by isolating $l$ clones from the evolutionary experiment, we can estimate the probability of sampling this adaptive event.

$$Prob(\text{sampling this adaptive event}) = 1-B( x=0, size=l, p = f )$$

$B( x=0, size=l, p = f )$ represents a binomial sampling process -- given the probability of sampling the adaptive event ($f$) and the number of clones sampled ($l$), the probability of missing this adaptive clone during sampling. We use this binomial sampling because the number of adaptive clones sampled is not important in the calculation. Instead, whether or not the adaptive clone is sampled is more important.

Combining these three steps together, we can calculate the probability to sample this particular adaptive event in one equation:

$Prob\ (sampling\ the\ adaptive\ event) = \int_0^t d\tau * s/\Gamma(N_b*\mu/c) * Exp[-N_b*\mu*(s/c)*\tau - Exp(-s*\tau)] * (1-B[x=0, size=l, p = c/(N_b*s) * Exp[s*(t-\tau)]])$.

**11.2 Parameters**

In our experiment, during each serial transfer cells are bottlenecked and then grown up for $T \approx 8$ generations. As described in the Supplementary Information in Levy, Blundell *et al*, the probability of the adaptive mutation arising during the first division of the cycle and surviving the bottleneck is $\approx N_b*u$. It is twice as likely to occur during the second division, but half as likely to survive the bottleneck, so the probability of a mutation entering at some point in the cycle and being present at a certain cell number once it is transferred to the next cycle is largely independent of where in the cycle it arises, always being proportional to $N_b*u$ with $N_b = 7E7$[1].

The variance ($2c$) in offspring number through the cycle (including Poisson noise of growth and bottleneck) was denoted as ~3.5 (Supplementary Information in Levy, Blundell *et al*[1]). Assuming Poisson noise each generation, the effective variance in offspring number *per generation* would be $2c = 3.5/T$[1].

Furthermore, ~90 neutral clones were isolated from Evo2D at generation 88 and sequenced in Venkataram, Dunn *et al*[3]. Based on the number of mutations in their genomes, we estimated a mutation rate ($\mu$) ~9E-10 per base-pair (bp) per division.

**11.3 Probability of not observing an optimal type with a 1bp mutational target size**

Assume that there is *1bp* in the yeast genome which can be mutated to the optimal type. We want to estimate the probability of sampling such an adaptive event from our evolutionary conditions. We consider the conservative scenario where the optimal type can only be generated by a specific nucleic acid change. Thus, the mutation rate $\mu = 9E\text{-}10/3 = 3E\text{-}10$ will be used for the following calculation.

Take the optimal type in Evo2D as an example. Based on its performance in fermentation and respiration, we can estimate its fitness per growth cycle in Evo2D (*s = 16h * fermentation performance + 28h * respiration performance = 2.56 per growth cycle*). As cells grow up to 8 generations per cycle, therefore, it is *fitness per generation* is *s/8*. Two replicates were conducted for Evo2D with *3,840* clones being isolated from replicate 1 and *960* clones being isolated from replicate 2. Using above parameters and equations, we can estimate that if such a 1bp mutational target size exists, the probabilities to sample this mutant from Evo2D replicate 1 and replicate 2 are *0.82* and *0.80*, respectively.

Similarly, the probabilities to observe an *1bp* optimal mutational event in Evo1D (*s = 16h * fermentation performance + 4h * respiration performance = 1.49 per growth cycle*) and Evo5D (*s = 40h * respiration performance + 60 * stationary phase performance = 2.98 per growth cycle*) are estimated. Note that the fitness of Evo5D optimal type is estimated by assuming a fermentation performance zero. As most adaptive clones do gain benefits from performance, the fitness of Evo5D optimal type is very likely to be under-estimated, providing a conservative calculation in our case. Two replicates were conducted for each evolutionary condition with *800* clones being isolated from each replicate. With a *1bp* mutational target size, the probability to sample this mutant from either Evo1D replicate 1 or replicate 2 is *0.39*; the probability to sample this mutant from either Evo5D replicate 1 or replicate 2 is *0.87*.

Based on above calculation, if the ancestral yeast is able to be mutated to maximize fermentation and respiration performances simultaneously by a single-step mutation, the probability of not observing such a mutant in both replicates of Evo1D and Evo2D will be *p = (1-0.39)*(1-0.39)*(1-0.82)*(1-0.8) = 0.013*.

Similarly, if the ancestral yeast is able to be mutated to maximize respiration and stationary phase performances by a single-step mutation, the probability of not observing such a mutant in both replicates of Evo5D will be *p = (1-0.87)*(1-0.87) = 0.017*.

Due to the difficulty in measuring fitness in Evo1/5D, the probability to not sample such optimal types is not calculated. If Evo1/5D was considered, the chance to miss such optimal types would be even smaller than what we estimated above.

**11.4 A conservative estimate of the above probability**

In our previous study (Li, Venkataram *et al*[2]), we noticed an inflation of fitness in one batch of fitness measurements; nonetheless, fitness measurements across batches are high correlated (Fig. 1). This batch of fitness measurements is used to infer fermentation and respiration performances among Evo2D adaptive clones in this study. Thus, the fitness estimate of the optimal type in the fermentation and respiration space may be affected and also inflated. Fitness measurements of Evo1D, Evo5D, and Evo1/5D adaptive clones were conducted independently in a different batch. Thus, these measurements likely don't suffer such an inflation.

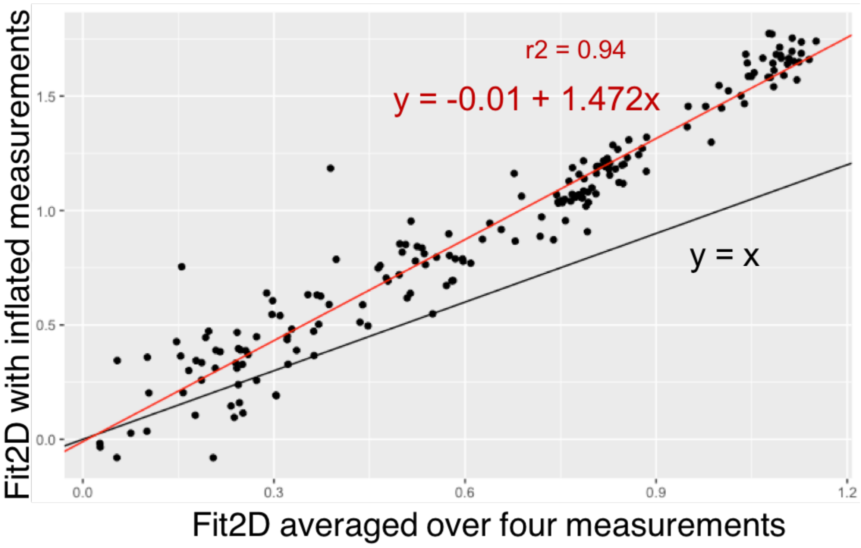


**Figure 1: Inflated fitness measurements among Evo2D adaptive clones.** Plot the batch of inflated fitness measurements against averaged fitness measurements across four batches under 2-day serial transfer condition (Fit2D). Each dot represents an Evo2D adaptive clone. The red line represents the linear fit between fitness measurements on x- and y-axis.

However, to have a conservative estimate, we corrected the estimated fitness of all optimal types by an inflation factor (*s_correct = (s + 0.01)/1.472*, Fig. 1) and generated corrected fitness estimates: *s per cycle* of Evo2D optimal type = *1.75*, s per cycle of Evo1D optimal type = *1.02*, and s per cycle of Evo5D optimal type = *2.04*.

With corrected fitness estimates, the probabilities to sample a single-mutation optimal type in Evo2D replicate 1 and replicate 2 are *0.6* and *0.54*, respectively. The probability to sample a single-mutation optimal type in either replicate of Evo1D is *0.08*. The probability to sample a single-mutation optimal type in either replicate of Evo5D is *0.65*.

Thus, if the ancestral yeast strain is able to be mutated to maximize fermentation and respiration performances simultaneously by a single-step mutation, the probability of not observing such a mutant in both replicates of Evo1D and Evo2D will be $p = (1-0.08)*(1-0.08)*(1-0.6)*(1-0.54) = 0.156$.

If the ancestral yeast is able to be mutated to maximize respiration and stationary phase performances by a single-step mutation, the probability of not observing such a mutant in both replicates of Evo5D will be $p = (1-0.65)*(1-0.65) = 0.123$, still strongly suggesting a lack of such a single-step mutation in the ancestral yeast genome.

#### 12 Viability Measurements

Two mutants carrying *FPK1* mutations, with one of them carrying an additional chromosome IX duplication, and two independent WT strains were cultured in monoculture for 5 days for viability measurements. The number of viable cells which formed colonies on plates was divided by the expected number measured by Coulter Counter to calculate viability -- the fraction of clones that are viable. Viability tests were conducted in two independent replicates with consistent results (Table S4).

#### Supplementary References

1. Levy, S. F. *et al.* Quantitative evolutionary dynamics using high-resolution lineage tracking. *Nature* **519**, 181–186 (2015).
2. Li, Y. *et al.* Hidden Complexity of Yeast Adaptation under Simple Evolutionary Conditions. *Curr. Biol.* **28**, 515-525.e6 (2018).
3. Venkataram, S. *et al.* Development of a Comprehensive Genotype-to-Fitness Map of Adaptation-Driving Mutations in Yeast. *Cell* **166**, 1585-1596.e22 (2016).
4. Zhao, L., Liu, Z., Levy, S. F. & Wu, S. Bartender: a fast and accurate clustering algorithm to count barcode reads. *Bioinformatics* **34**, 739–747 (2018).
5. Kryazhimskiy, S., Rice, D. P., Jerison, E. R. & Desai, M. M. Global epistasis makes adaptation predictable despite sequence-level stochasticity. *Science* **344**, 1519–1522 (2014).
6. Freed, D. N., Aldana, R., Weber, J. A. & Edwards, J. S. The Sentieon Genomics Tools - A fast and accurate solution to variant calling from next-generation sequence data. *bioRxiv* 115717 (2017). doi:10.1101/115717
7. Cingolani, P. *et al.* A program for annotating and predicting the effects of single nucleotide polymorphisms, SnpEff. *Fly (Austin)* **6**, 80–92 (2012).
